# The PD-L1 and TLR7 dual-targeting nanobody-drug conjugate exerts potent antitumor efficacy by orchestrating innate and adaptive immune responses

**DOI:** 10.1101/2021.04.11.439388

**Authors:** Xiaolu Yu, Yiru Long, Binfan Chen, Yongliang Tong, Xiaomin Jia, Ji Zhou, Feng Tang, Pan Xu, Yuhan Cao, Wei Huang, Jin Ren, Yakun Wan, Jianhua Sun, Guangyi Jin, Likun Gong

## Abstract

A variety of tumors are insensitive to immune checkpoint blockade (ICB) therapy. We propose that ICB therapy alone is insufficient to fully reactivate antitumor T cells, while effective mobilization of antigen-presenting cells (APCs) to assist adaptive immune cell activation can lead to potent antitumor effects with broad responsiveness. The Toll-like receptor 7 (TLR7) agonist SZU-101 we developed can induce the innate immune response against tumors and increase the immunogenicity of tumors. Interestingly, SZU-101-induced upregulation of programmed death ligand 1 (PD-L1) expression in tumor tissues can further enhance the response rate of the PD-L1 antibody. In addition, PD-L1 nanobodies have better solid tumor penetration ability, and because of this ability, they can be used to precisely deliver SZU-101 to tumor tissues. Therefore, a PD-L1 and TLR7 dual-targeting nanobody-drug conjugate (NDC), a novel drug molecule, was developed. We found that TLR7 agonists and PD-L1 nanobodies act synergistically and that NDC treatment reshapes the tumor immune microenvironment, activates both innate and adaptive immune cells, and exerts antitumor effects in both “hot” and “cold” tumors primarily through CD8^+^ T cells and natural killer (NK) cells. Our data show that a PD-L1 and TLR7 dual-targeting NDC can exhibit potent antitumor efficacy by orchestrating innate and adaptive immune responses and shows good prospects for clinical development.

**One Sentence Summary:** Based on results showing that TLR7 agonists and PD-L1 nanobodies exert synergistic antitumor effects, a PD-L1 and TLR7 dual-targeting nanobody-drug conjugate that we developed shows good prospects for clinical development because it can orchestrate innate and adaptive immune responses, reshape the tumor immune microenvironment, and exert potent antitumor effects against both “hot” and “cold” tumors.

## INTRODUCTION

In the tumor microenvironment (TME), programmed death ligand 1 (PD-L1) molecules, which are mainly expressed on tumor cells, macrophages and dendritic cells (DCs), interact with programmed death 1 (PD-1) on T cells, leading to antitumor immunity inhibition and T cell exhaustion, which are key mechanisms of tumor immune escape (1–3). PD-1/PD-L1 immune checkpoint blockade therapy (ICB) is effective against many cancer types (4–9), but only a small fraction of patients responds to treatment (10). Determining the tumor resistance mechanisms for PD-1/PD-L1 antibody therapy and improving therapeutic efficacy are urgent areas to be addressed (11). PD-1/PD-L1 antibodies are being combined with diverse targeted drugs to increase treatment response rates in preclinical and clinical trials (12, 13).

PD-1/PD-L1 antibody therapy alone is more efficacious for highly immunogenic “hot” tumors with adequate T cell infiltration and/or high PD-L1 expression, such as melanoma and non-small cell lung cancer, and less efficacious for poorly immunogenic “cold” tumors with inadequate T cell infiltration and/or low PD-L1 expression, such as breast cancer and pancreatic cancer (14, 15). Thus, the combination of drugs that facilitate antigen presentation and PD-1/PD-L1 antibodies can mobilize both innate and adaptive immunity to break tumor immune tolerance, increase tumor immunogenicity, and transform “cold” tumors into “hot” tumors, for which a broadly responsive therapeutic effect can be expected (*16–18*).

Toll-like receptor 7 (TLR7), a target for immunogenicity promotion and immune enhancement, is predominantly expressed in macrophages, DCs, natural killer (NK) cells and B cells (19, 20). Our studies and those of others have shown that TLR7 agonists can inhibit tumor growth in a wide range of tumor models (*21–27*). By stimulating TLR7, antigen-presenting cells (APCs) can be activated, and subsequently, their antigen-presenting function is enhanced to facilitate T cell activation (*28, 29*). In addition, the increased intratumoral infiltration of NK cells, cytotoxic T cells and antigen-specific interferon (IFN)-secreting effector cells is one of the antitumor mechanisms of TLR7 agonists (*30–32*). The use of a TLR7 agonist is a potential option to improve the response rates of ICB therapy (*33*).

In addition to the need to improve the immunogenicity of tumors, favorable tumor penetration of drug molecules is also crucial. Solid tumors have a dense extracellular matrix (ECM) that makes it difficult for conventional monoclonal antibodies to penetrate deeply into the tumor interior (*34*). Nanobodies have good tumor penetration and specificity, making them suitable for targeted therapy and drug delivery (*35*). Based on nanobodies, nanobody-drug conjugates (NDCs) are new forms of antibody-drug conjugates (ADCs) (*36*). Many NDCs that couple diverse immunotoxins or drugs with nanobodies have been developed and have shown beneficial antitumor effects in a diversity of tumor models (*37*). Our study and those of others have shown that PD-L1 antibodies can be used to deliver drugs to solid tumors (*17, 38*), and PD-L1 nanobodies are promising constituents for use in the development of NDCs.

We developed a PD-L1 and TLR7 dual-targeting NDC, which has good tumor permeability, enables precise delivery of TLR7 agonists to intratumoral target cells and attenuates the intratumoral immune tolerance environment. The PD-L1 and TLR7 dual-targeting NDC represents a combination therapeutic strategy that boosts antigen presentation and breaks immune tolerance, leading to the integration of innate and adaptive immunity for potent antitumor effects and broad responsiveness.

## RESULTS

### Screening and identification of PD-L1 nanobodies

Compared to monoclonal antibodies, nanobodies have many advantages, such as small size, high stability, and low immunogenicity (*39*). After preparation of the PD-L1 antigen and immunization of a camel, we constructed a high-quality phage display nanobody library with a capacity of 1.8 × 10^6^ colony forming units (CFU) and a 96% insert rate of the antibody gene (fig. S1), from which nanobody candidates were screened and confirmed to block PD-1/PD-L1 binding (fig. S2, A and B). To endow antibodies with antibody-dependent cellular cytotoxicity (ADCC) effects in vivo, candidates were linked to Fc fragments to form fusion proteins, and the activity of these fusion proteins was evaluated in vitro. The optimized candidate (termed Nb16) was selected (fig. S2C). Sodium dodecyl sulfate polyacrylamide gel electrophoresis (SDS-PAGE) results showed that the purified proteins had a purity >95% (fig. S2D). Nb16 showed a strong binding affinity for PD-L1, with a 50% effective concentration (EC50) value of 0.04336 μg/mL or 0.516 nM (Fig. 1A). Nb16 strongly blocked the interaction between PD-1 and PD-L1, with a 50% inhibitory concentration (IC50) value of 0.5691 μg/mL or 6.78 nM (Fig. 1B). The binding and blocking activity of Nb16 was verified in HEK 293T cells stably expressing PD-L1 (Fig. 1C and D).

**Fig. 1.**
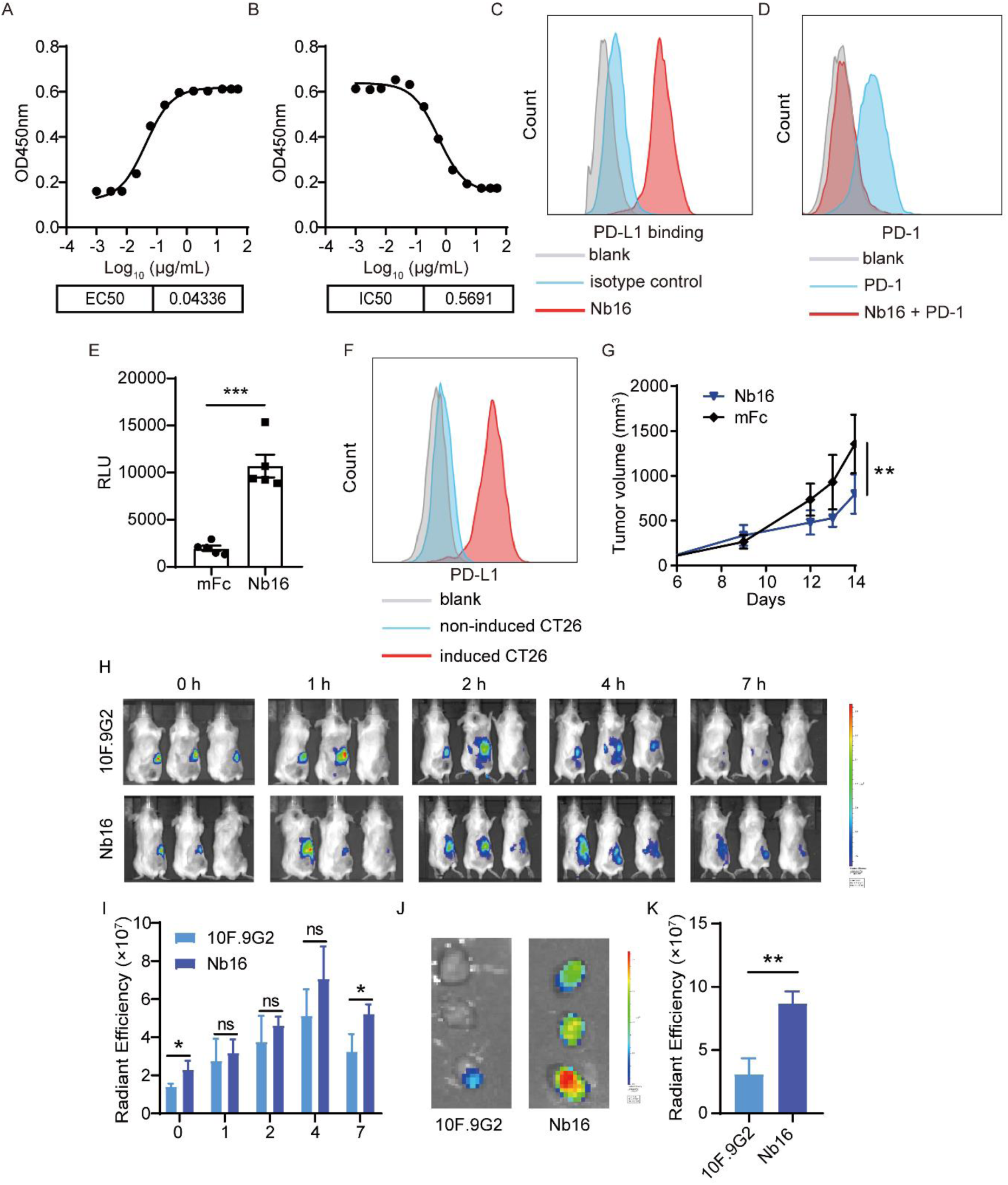
The PD-L1 nanobody inhibits tumor growth and has better solid tumor penetration. (**A**) Binding affinity of Nb16 to PD-L1. Serial dilutions of Nb16 series were incubated with plate-bound PD-L1, and binding was assessed via anti–Fc ELISA. (**B**) PD-1/PD-L1 blocking activity of Nb16. Serial dilutions of Nb16 were incubated with PD-1-biotin (10 μg/mL) and plate-bound PD-L1, and PD-1 binding was assessed via streptavidin ELISA. (**C)** Flow cytometry histograms showing the binding of Nb16 to PD-L1 on HEK 293T cells stably expressing PD-L1. (**D**) Flow cytometry histograms showing that Nb16 blocked the PD-1/PD-L1 interaction in HEK 293T cells stably expressing PD-L1. (**E**) Effect of Nb16 on blocking the inhibitory signal from CT26 cells (induced by IFN-γ) to Jurkat-NFAT-mPD-1-luciferase cells. The data are shown in relative luciferase units (RLU). (**F**) Flow cytometry histograms showing the high expression of PD-L1 in CT26 cells induced by IFN-γ (100 ng/mL) for 24 h. (**G**) BALB/c mice were subcutaneously inoculated with 1×10^6^ of CT26 cells (induced by IFN-γ). Mice bearing tumors were treated with 200 µg of Nb16 or mFc every three days. The tumor growth in mice is shown (n = 6). (**H**) In vivo imaging showed the enhanced tumor accumulation of Nb16. (**I**) In vivo radiant efficiency at the tumor sites. (**J**) Ex vivo imaging of the tumors. (**K**) Ex vivo radiant efficiency of the tumors. The data are presented as the mean ± SEM and are representative of at least two independent experiments. * p < 0.05; ** p < 0.01; ns, not significant as determined by unpaired t test.

### Antitumor activity and solid tumor penetration of Nb16

Reversed PD-L1 repression can be measured by the level of nuclear factor of activated T cell (NFAT) activation in response to PD-L1 antibody treatment (*40*). Thus, we constructed Jurkat-NFAT-mPD-1-luciferase cells to perform a reporter assay (fig. S3). Nb16 blocked the inhibitory signal from CT26 cells (induced by IFN-γ to express high levels of PD-L1) to T cells (Fig. 1, E and F). Subsequently, the in vivo activity of Nb16 was assessed. CT26 tumors (induced by IFN-γ) were treated with Nb16 twice a week. The results showed that Nb16 treatment significantly suppressed the growth of CT26 tumors with a tumor inhibition rate of 39.1% (Fig. 1G).

To evaluate the tumor penetration ability and tumor targeting delivery efficiency of Nb16, Nb16 was labeled with fluorochrome Cy7 and then injected into mice to detect the in vivo distribution from 0 h to 7 h. PD-L1 monoclonal antibody 10F.9G2 was used as a comparison. Fluorescence at the tumor site was visible within 2 h (Fig. 1H). The fluorescence intensity in the tumor gradually increased and peaked from 4 to 7 h (Fig. 1I). Drug tumor accumulation was significantly higher in mice receiving Nb16, and the ex vivo imaging of dissected tumor tissue also showed 2.7-fold higher tumor accumulation in the Nb16 group than in the 10F.9G2 group (Fig. 1, J and K). This demonstrated that the PD-L1 nanobody had enhanced solid tumor penetration compared with PD-L1 monoclonal antibodies. In summary, the PD-L1 nanobody Nb16 with prominent antitumor activity and tumor penetration has potential for use in antitumor drug delivery.

### The association of TLR7 expression with the prognosis and immune infiltration of TCGA cohorts

Studies have shown that TLR7 expression levels are low in many tumors (*41–43*). We characterized the expression pattern of TLR7 in different tumors and also found many of them with low TLR7 expression through TIMER2.0 (*44*) based on The Cancer Genome Atlas (TCGA) datasets (Fig. 2A). Cancer patients with low TLR7 expression have a remarkably shorter overall survival (OS) (Fig. 2, B-D). Impaired TLR7 signaling may be one of the mechanisms for tumor immune escape. In contrast, studies have shown that TLR7 is highly expressed in some tumors and shows tumorigenic effects, which may be related to long-term low-level activation of TLR7 by endogenous weak agonists in the TME (*45–47*).

**Fig. 2.**
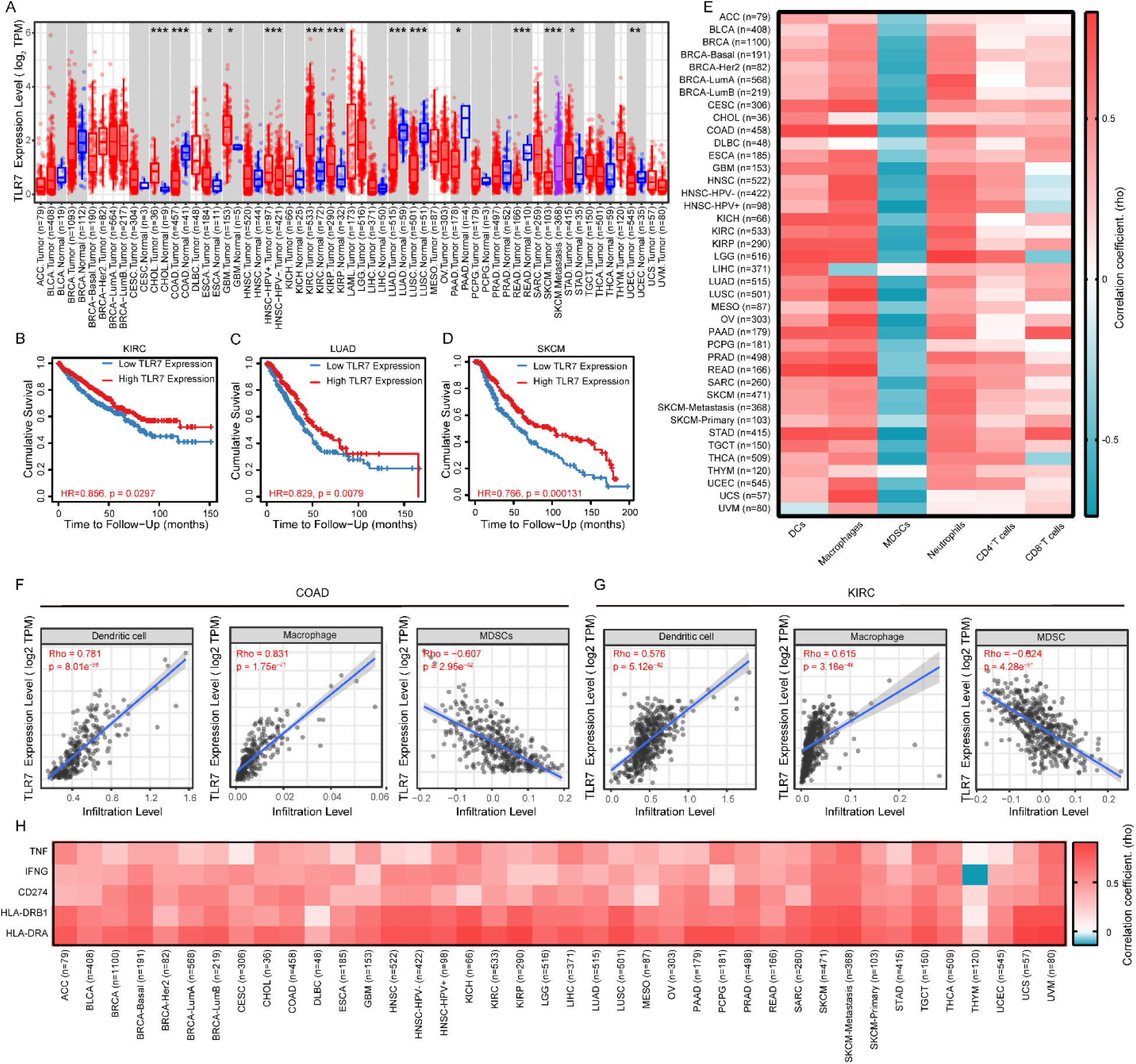
TLR7 expression correlates with prognosis and immune infiltration in TCGA cohorts. (**A**) Human TLR7 expression levels in different tumor types from the TCGA database were determined by TIMER2.0 (*P < 0.05, **P < 0.01, ***P < 0.001 by the Wilcoxon test). (**B** to **D**) Kaplan-Meier curves showing the OS in patients with different tumor types as indicated in the TCGA database and determined by TIMER2.0. (**E** to **G**) The correlation of TLR7 expression with immune infiltration level in different tumor types as indicated in the TCGA database was determined by TIMER2.0. Heat map showing the correlation between TLR7 expression levels and immune cell (DC, macrophage, MDSC, neutrophil, CD4^+^ T cell and CD8^+^ T cell) infiltration levels in different tumor types (E). Scatter plots showing the correlation between TLR7 expression levels and immune cells (DCs, macrophages and MDSCs) in COAD (F) and KIRC (G). (**H**) Correlation analysis of TLR7 and related genes (TNF, IFNG, CD274, HLA-DRB1 and HLA-DRA) in different tumor types in the TCGA database as determined by TIMER2.0. TIMER2.0, Tumor Immune Estimation Resource.

Furthermore, TLR7 levels showed a strong positive correlation with intratumoral infiltration of innate immune cells (macrophages, DCs and neutrophils) and a strong negative correlation with intratumoral infiltration of myeloid-derived suppressor cells (MDSCs) (Fig. 2, E-G). In contrast, the expression level of TLR7 and the intratumoral infiltration of adaptive immune CD8^+^ T cells and CD4^+^ T cells did not show a significant correlation (Fig. 2E).

On the other hand, the expression levels of TLR7 with tumor necrosis factor-α (TNF-α), IFN-γ, and major histocompatibility complex class II (MHC II) in tumors showed a strong positive correlation (Fig. 2H and fig. S4), suggesting that activation of TLR7 can promote the activity of antigen-presenting machinery and the secretion of the antitumor cytokines. Notably, tumors with high levels of TLR7 were also accompanied by high expression of PD-L1 (Fig. 2H and fig. S4), suggesting that the combination of TLR7 agonists with the PD-L1 antibodies may be a rational strategy for cancer therapy.

### Activation of innate immunity by the TLR7 small-molecule agonist SZU-101

We developed SZU-101 (*23*), which is a potent TLR7 small-molecule agonist (Fig. 3A). SZU-101 promoted splenocyte proliferation in a dose-dependent manner in vitro, showing a prominent immune activating effect (Fig. 3B). Furthermore, SZU-101 can activate mouse bone marrow-derived DCs (BMDCs), stimulate the expression of their maturation markers CD80 and CD86, and enhance the expression of MHC II (Fig. 3, C-E). Thus, SZU-101 can effectively promote the maturation of DCs and enhance their antigen-presenting ability. Considering that tumor-associated macrophages (TAMs) are among the important factors affecting tumor progression (*48*), we further tested the effect of SZU-101 on bone marrow-derived macrophages (BMDMs). After macrophages were induced to differentiate into the M2 type, we found that SZU-101 repolarizes macrophages. SZU-101 significantly upregulated the M1 macrophage marker CD86, significantly downregulated the M2 macrophage marker CD206, and simultaneously stimulated macrophages to express the antitumor cytokine IFN-γ (Fig. 3, F-H). In addition, SZU-101 stimulated NK cells to highly express the activation markers CD69 and CD25 (Fig. 3, I and J). These results suggest that SZU-101 can activate innate immunity and has potential antitumor activity.

**Fig. 3.**
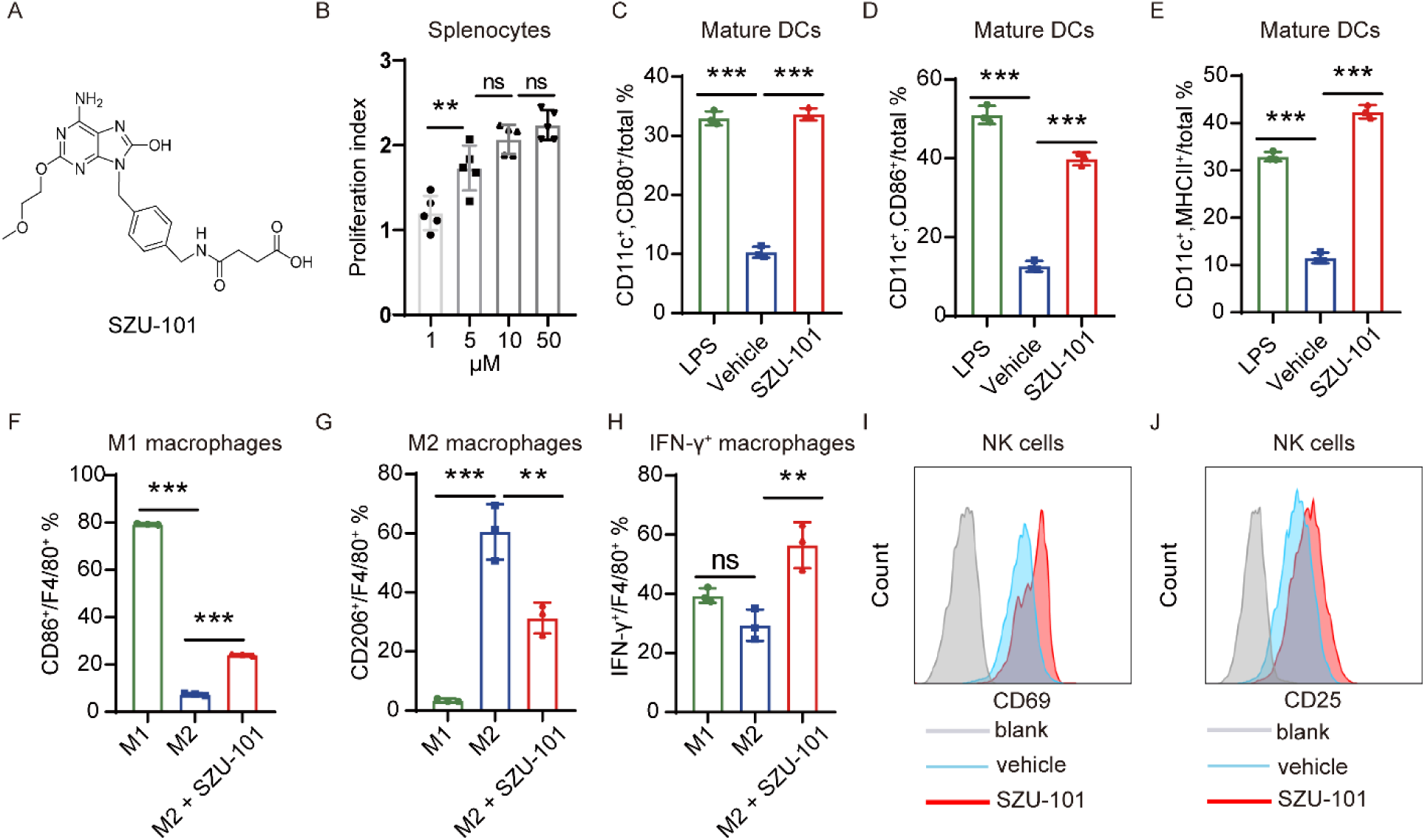
Activation of innate immunity by TLR7 agonist SZU-101. (**A**) The structure of SZU-101. (**B**) Spleen cells obtained from BALB/c mice were treated with either complete medium or SZU-101 (1, 5, 10 or 50 μM, n = 5). An MTT assay was performed to determine the cell proliferation activity, shown as the proliferation index, which is calculated as the fold change with respect to the unstimulated control cultures. (**C** to **E**) BALB/c mouse bone marrow-derived dendritic cells (BMDCs) were treated with complete medium, LPS or 50 μM of SZU-101 for 24 h (n = 3). CD80 (C), CD86 (D) and MHC II (E) on DCs were detected by flow cytometry. (**F** to **H**) BALB/c mouse bone marrow-derived macrophages (BMDMs) induced to differentiate into M2 macrophages were stimulated with 50 μM of SZU-101 for 24 h (n = 3). CD86 (F), CD206 (G) and IFN-γ (H) expression in macrophages was determined by flow cytometry. (**I** and **J**) Flow cytometry histograms showing the high expression of CD69 (I) and CD25 (J) on NK92 cells stimulated with 50 μM of SZU-101 for 24 h. The data are presented as the mean ± SEM and are representative of at least two independent experiments. ** p < 0.01; *** p < 0.001; ns, not significant as determined by one-way ANOVA with multiple comparison tests.

### SZU-101 increased PD-L1 expression in tumors, and intratumoral administration is required

SZU-101 promoted the secretion of the antitumor cytokines TNF-α and IFN-γ from splenocytes in a dose-dependent manner in vitro, which is consistent with the analysis of clinical TCGA data, suggesting a prominent induction effect of the antitumor immune response (Fig. 4, A and B). Considering that TNF-α and IFN-γ are potent inducers of PD-L1 and that a clinical TCGA data analysis showed a significant positive correlation between TLR7 expression levels and PD-L1, we investigated the effect of SZU-101 on PD-L1 expression levels in CT26 tumor cells, DCs and macrophages in vitro. SZU-101 significantly increased PD-L1 expression on DCs and macrophages in the splenocyte population (Fig. 4, C and D). In addition, in the coculture system of CT26 cells and splenocytes, SZU-101 induced high PD-L1 expression in CT26 tumor cells (Fig. 4E), while SZU-101 did not directly promote PD-L1 expression in CT26 cells cultured alone (Fig. 4F).

**Fig. 4.**
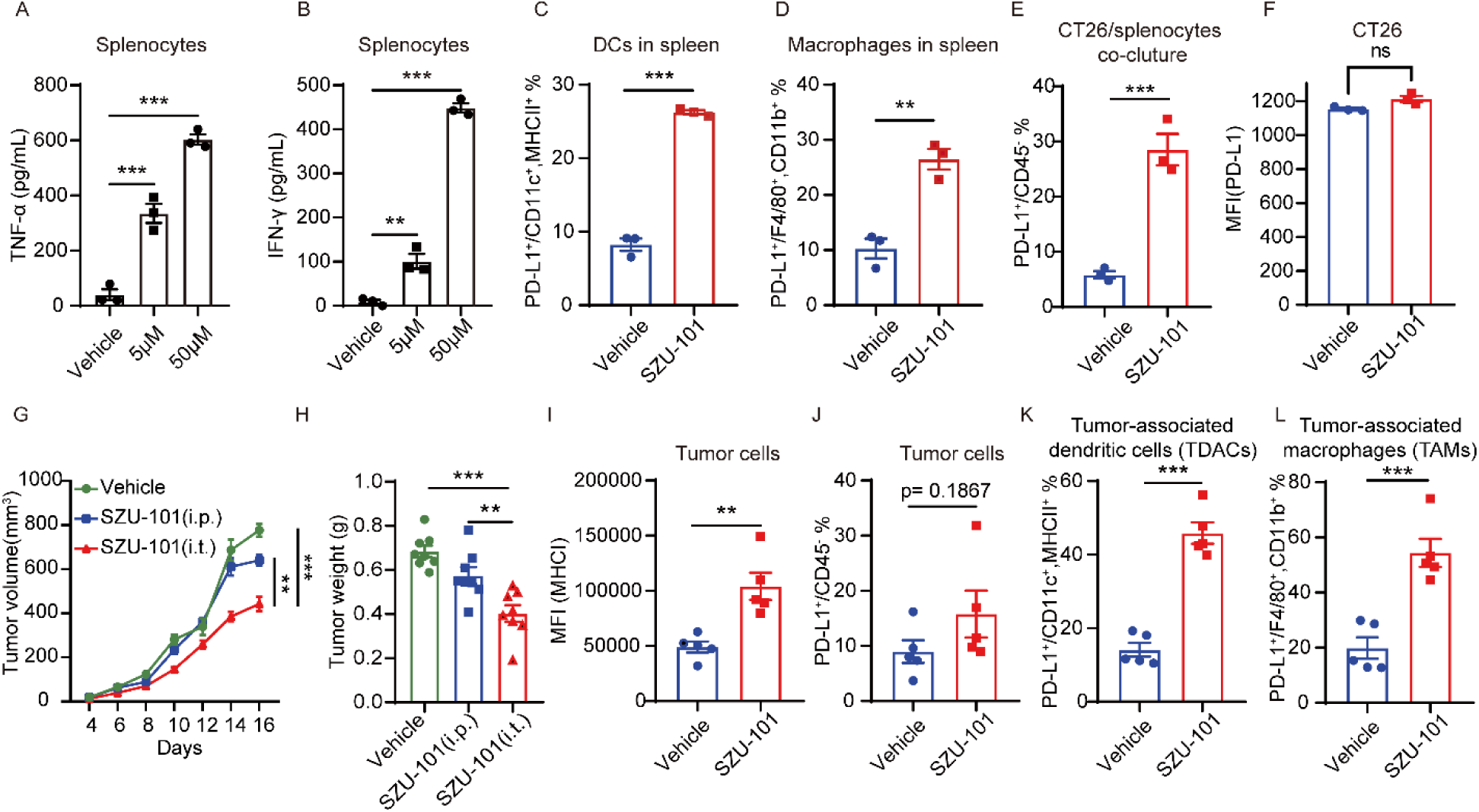
The TLR7 agonist SZU-101 induced the upregulation of intratumoral PD-L1 levels and required intratumoral administration. (**A** and **B**) Spleen cells obtained from BALB/c mice were treated with PBS or SZU-101 (5 μM or 50 μM, n = 3). TNF-α (A) and IFN-γ (B) secreted by spleen cells were detected by anti-TNF-α and anti-IFN-γ ELISAs. (**C** and **D**) Spleen cells obtained from BALB/c mice were treated with 50 μM SZU-101 for 24 h (n = 3). PD-L1 expression on DCs (C) and macrophages (D) was determined by flow cytometry. (**E**) CT26 cells were cocultured with spleen cells obtained from BALB/c mice (n = 3). After treatment with 50 μM of SZU-101 for 24 h, PD-L1 expression on CT26 cells was determined by flow cytometry. (**F**) Flow cytometry histograms showing the PD-L1 levels on CT26 cells induced by 50 μM of SZU-101 or a vehicle. (**G** and **H**) BALB/c mice were subcutaneously inoculated with 1×10^6^ of CT26 cells. Mice bearing tumors were treated with SZU-101 (3 mg/kg) intratumorally (i.t.) and intraperitoneally (i.p.) or with a vehicle every two days. Tumor growth in mice is shown (n = 8). (**I**) MHC I expression on CD45- cells in CT26 tumor tissues from tumor-bearing BALB/c mice treated with SZU-101 or a vehicle was determined by flow cytometry (n = 5) MFI, median fluorescence intensity. (**J** to **L**) PD-L1 expression on CD45- cells, DCs, and macrophages in CT26 tumor tissues from tumor-bearing BALB/c mice treated with SZU-101 or a vehicle was determined by flow cytometry (n = 5). The data are presented as the mean ± SEM and are representative of at least two independent experiments. ** p < 0.01; *** p < 0.001; ns not significant by one-way ANOVA with multiple comparison tests or unpaired t test.

Our previous study showed that SZU-101 significantly inhibited the growth of EL4 and B16 tumors (*23, 27*). Similarly, SZU-101 significantly inhibited CT26 tumor growth (Fig. 4, G and H). However, systemic administration of SZU-101 did not induce antitumor activity to the extent induced by local intratumoral administration (Fig. 4, G and H), which suggested that precise delivery of SZU-101 to tumors can achieve better antitumor effects. Moreover, SZU-101 promoted the expression of major histocompatibility complex class I (MHC I) on tumor cells, suggesting that SZU-101 can enhance the immunogenicity of tumors (Fig. 4I). Notably, tumor cells and antigen-presenting cells (DCs and macrophages) were also induced to overexpress PD-L1 by SZU-101 in vivo (Fig. 4, J-L), suggesting that combined PD-L1 antibody and TLR7 agonist therapy may achieve better antitumor efficacy.

### The PD-L1 nanobody and TLR7 agonist exert synergistic antitumor effects

SZU-101 can activate innate immunity, and Nb16 can restore adaptive immunity; therefore, we tested the effect of SZU-101 and Nb16 combination treatment (Fig. 5A). The growth of CT26 tumors treated with a combination of Nb16 and SZU-101 was significantly inhibited compared to single-agent treatment (Fig. 5, B and C and fig. S5A). Thus, Nb16 and SZU-101 exert synergistic antitumor effects. In addition, the body weight curve of mice after administration suggested good safety of the combination treatment (fig. S5B).

**Fig. 5.**
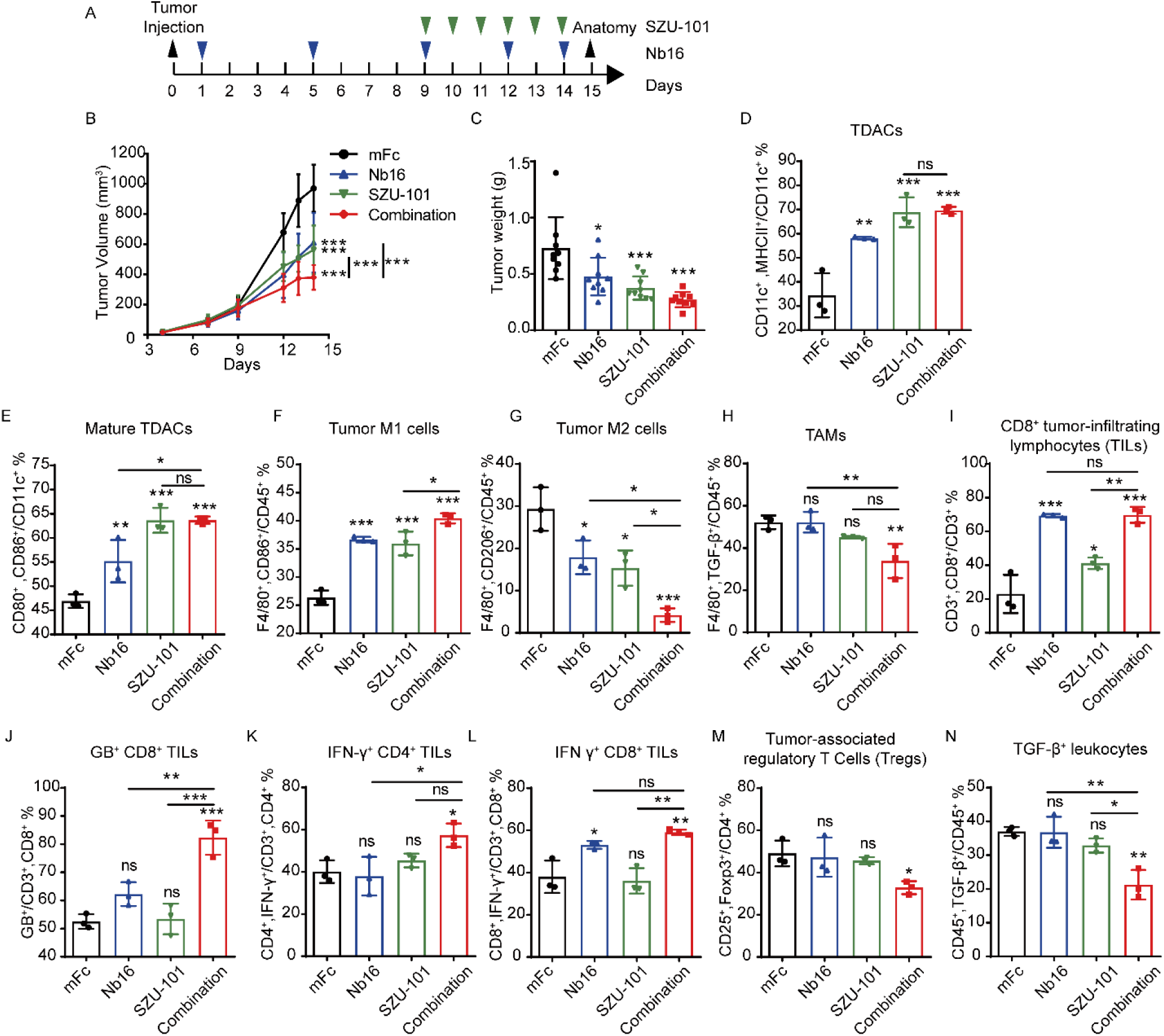
The PD-L1 nanobody and TLR7 small molecule agonist exert synergistic antitumor effects. (**A**) Treatment scheme. BALB/c mice were subcutaneously inoculated with 1×10^6^ of CT26 cells (induced by IFN-γ). Mice bearing tumors were treated with 200 μg of mFc, SZU-101 (3 mg/kg), 200 μg of Nb16 or a combination of these compounds. (**B**) Tumor growth in mice treated as described in the scheme (n = 9). (**C**) The weight of the tumors from mice treated as described in the scheme is shown (n = 9). (**D** to **N**) Immune effector cells in tumors from mice treated as described in the scheme were quantified by flow cytometry (n = 3). Scatter plots show the proportion of cells in a specific cell population. The percentages of MHC II^+^ DCs (D), mature DCs (E), M1 macrophages (F), M2 macrophages (G), TGF-β^+^ macrophages (H), CD8^+^ T cells (I), GB^+^ CD8^+^ T cells (J), IFN-γ^+^ CD4^+^ T cells (K), IFN-γ^+^ CD8^+^ T cells (L), Tregs (M) and TGF-β^+^ leukocytes (N) are shown based on their respective markers. The data are presented the mean ± SEM and are representative of at least two independent experiments. * p < 0.05; ** p < 0.01; *** p < 0.001; ns, not significant as determined by one-way ANOVA with multiple comparison tests.

Next, we used flow cytometry to quantify immune effector cells in mFc-treated, Nb16-treated, SZU-101-treated and combination-treated tumors. SZU-101 treatment and combination treatment significantly increased DCs infiltration and activation in the TME to the same extent (Fig. 5, D and E and fig. S6A-B), indicating that the changes of DCs in the combination group were mainly caused by SZU-101. Combination therapy further increased infiltration of M1 macrophages (Fig. 5F and fig. S6C) and decreased infiltration of M2 macrophages (Fig. 5G and fig. S6D) compared to single agents. And only the combination treatment significantly downregulated transforming growth factor-β (TGF-β) expression in macrophages (Fig. 5H and fig. S6E). Further analysis of tumor-infiltrating lymphocyte alterations showed that both the number of regulatory T cells (Tregs) and TGF-β-associated immune cells were significantly decreased after treatment with combination therapy, whereas neither was affected by administration of Nb16 or SZU-101 (Fig. 5, M and N and fig. S6J-K). Thus, the combination treatment can attenuate the immunosuppressive effect of the TME. Further analysis of the activation of adaptive immunity showed that Nb16 treatment and the combination treatment increased the infiltration of intratumoral CD8^+^ T cells equally, while SZU-101 did not, suggesting that the changes in CD8^+^ T cells in the combination group were mainly caused by Nb16 (Fig. 5I and fig. S6F). Notably, only the combination treatment promoted the expression of granzyme B (GB) and IFN-γ in CD8^+^ T cells and IFN-γ in CD4^+^ T cells, showing elevated T cell antitumor activity (Fig. 5, J-L and fig. S6, G-I). In brief, Nb16 combined with SZU-101 can synergistically control tumor growth by reshaping the TME, relieving tumor immunosuppression and activating antitumor immune responses.

### Combination therapy with the PD-L1 nanobody and TLR7 agonist promotes the expression of immune cell signature genes

To examine the gene expression changes induced by the combined Nb16 and SZU-101 treatment, we analyzed CT26 tumor tissues by RNA sequencing (RNA-seq) (fig. S7A). Compared to isotype control treatment, combination treatment resulted in differential gene expression, which were mainly enriched in immune system-related pathways, particularly antigen presentation and cytotoxic cell function pathways.(fig. S7B).

To further assess the effects of combination treatment on immune infiltration and immune processes, we analyzed the expression of gene signatures associated with distinct immune cell types, IFN responses, antigen presentation processes and apoptosis. The signature genes and analysis methods were described in published literature (*49, 50*). The combination treatment increased the signature scores of T cells, NK cells, DCs, the IFN-α response, the IFN-γ response, the antigen presentation process and apoptosis compared to mFc and the single agents (Fig. 6, A-H). Since T cells have functionally complex subpopulations, we used ImmuCellAI to analyze T cell infiltration within tumor tissues (*51*). The results showed that the combination therapy caused a reduction in the percentage of naive T cells in the tumor (Fig. 6I). Moreover, the combination treatment led to an increase in intratumoral Th1 cells and a decrease in Th17 cells (Fig. 6I). In particular, the combination treatment significantly increased the intratumoral infiltration of cytotoxic cells such as NK cells and CD8^+^ T cells (Fig. 6I).

**Fig. 6.**
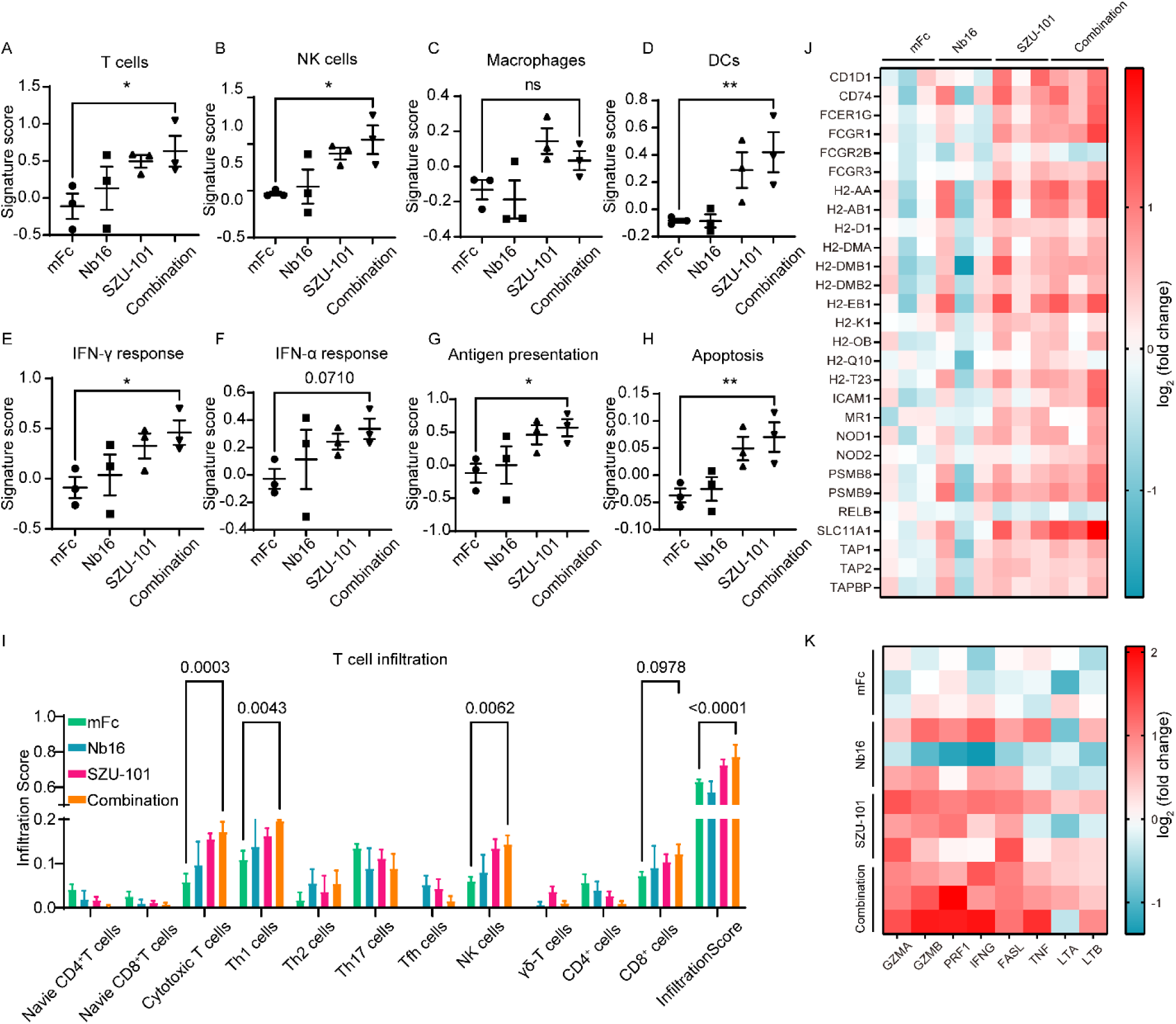
The Nb16 and SZU-101 combination treatment promotes gene expression associated with innate and adaptive immune cells. RNA-sequencing analysis of tumor tissue from BALB/c mice (n = 3 mice per group) that were bearing 1×10^6^ of CT26 cells (induced by IFN-γ) and treated with 200 μg of mFc, SZU-101 (3 mg/kg), 200 μg of Nb16 or a combination of these compounds. (**A** to **H**) Gene expression signatures associated with T cells (A), NK cells (B), macrophages (C), DCs (D), the IFN-γ response (E), the IFN-α response (F), the antigen presentation process (G) and apoptosis (H). Signature scores [defined as the mean log_2_(fold change) among all genes in the signature] are shown as scatter plots. (**I**) T cell infiltration analysis by ImmuCellAI. (**J**) Heat map shows that antigen presentation process–associated genes changed after treatment [defined as the log_2_(fold change)]. (**K**) Heat map showing that cytotoxic cell effector molecule– associated genes changed after treatment [defined as the log_2_(fold change)]. The data are presented as the means ± SEM. * p < 0.05; ** p < 0.01; ns, not significant as determined by one-way ANOVA with multiple comparison tests.

This analysis suggested that Nb16 and SZU-101 treatment may affect the infiltration and function of APCs and cytotoxic cells; therefore, we further analyzed the expression levels of genes characterizing the antigen presentation process and cytotoxic cell effector molecules. The combination therapy significantly increased the activity of intratumoral antigen-presenting machinery and promoted the expression of effector molecules of cytotoxic cells, such as IFN-γ, TNF-α, granzyme B, perforin, and FASL (Fig. 6, J and K). The analysis of the RNA-seq results revealed that Nb16 and SZU-101 show synergistic action, and SZU-101 treatment notably promoted the antigen presentation process in vivo and facilitated the activation and function of T cells, which was consistent with our aforementioned findings.

In summary, the RNA-seq results suggested that the combination of the PD-L1 nanobody and TLR7 agonist exerted potent antitumor efficacy through promoting antitumor cell infiltration, antigen presentation and cytotoxic effects.

### Preparation and identification of the PD-L1 and TLR7 dual-targeting nanobody-drug conjugate

Based on the good antitumor effect of Nb16 and SZU-101 combination therapy, considering that PD-L1 nanobody possesses effective tumor penetration and targeting properties and SZU-101 promotes intratumoral PD-L1 expression, coupling SZU-101 with Nb16 to prepare a dual-targeting NDC was thought to enable a better distribution and localization of both compounds in tumor tissues for better antitumor efficacy.

To prepare the dual-targeting NDC, SZU-101 was connected to PEG to yield the intermediate compound SZU-107 (fig. S8A). Comparing the in vitro immune cell activation effect of SZU- 101 and SZU-107, we found that both had equal effects in terms of activating innate immunity, indicating that PEGylation did not affect the biological activity of SZU-101 (fig. S8, B-H). The novel dual-targeting NDC Nb16-SZU-101 was formed by coupling carboxyl group-containing SZU-107 and primary amine group-containing Nb16 via the condensation agent EDC (Fig. 7A). The compounds were identified by mass spectrometry before and after coupling, and the drug-to-antibody ratio (DAR) of the NDC was determined to be 4.5 according to the change in the molecular weight of the antibody (fig. S8, I and J). To determine whether the activity of the nanobodies was affected before and after coupling, the EC50 and IC50 values of the antibodies were measured before and after coupling. The results showed no significant difference in the EC50 and IC50 values of the antibodies before and after coupling, indicating that NDC retained the target binding and blocking activities of the original nanobody (Fig. 7, B and C). As a result, we obtained a novel PD-L1 and TLR7 dual-targeting NDC.

**Fig. 7.**
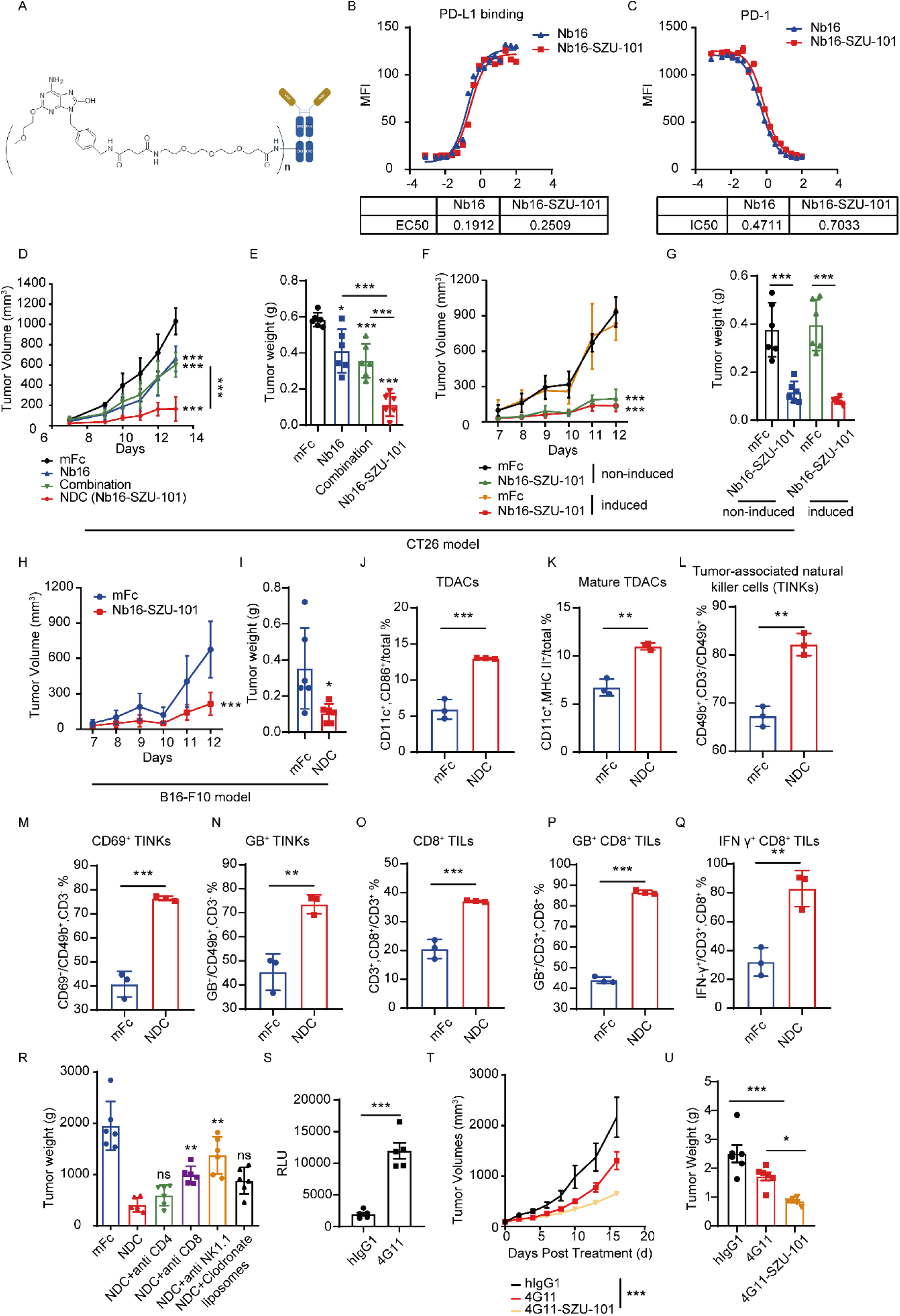
The PD-L1 and TLR7 dual-targeting NDC plays an antitumor effect by combining innate and adaptive immunity. (**A**) The structure schematic of Nb16-SZU-101. (**B**) Binding affinity of Nb16 and Nb16-SZU-101 for PD-L1. Serial dilutions of Nb16 and Nb16-SZU-101 were incubated with HEK 293T cells stably expressing PD-L1, and binding was assessed via flow cytometry. MFI, median fluorescence intensity. (**C**) PD-1/PD-L1 blocking activity of Nb16 and Nb16-SZU-101. Serial dilutions of Nb16 and Nb16-SZU-101 were incubated with HEK 293T cells stably expressing PD-L1, 10 μg/mL of PD-1-biotin and PD-L1. PD-1 binding was assessed via flow cytometry. MFI, median fluorescence intensity. (**D**) BALB/c mice were subcutaneously inoculated with 1×10^6^ of CT26 cells (induced by IFN-γ). Mice bearing tumors were treated with 200 μg of mFc, 200 μg of Nb16, combination or 200 μg of NDC. Tumor growth in mice is shown (n = 6). (**E**) The weight of tumors from mice is shown (n = 6). (**F** and **G**) BALB/c mice were subcutaneously inoculated with 1×10^6^ of CT26 cells (induced by IFN-γ or not). Mice bearing tumors were treated with 200 μg of mFc or 200 μg of NDC. Tumor growth (F) in mice and the weight of tumors (G) are shown (n = 6). (**H** and **I**) C57BL/6 mice were subcutaneously inoculated with 1×10^6^ of B16-F10 cells. Mice bearing tumors were treated with 200 μg of mFc or 200 μg of NDC. Tumor growth (H) in mice and the weight of tumors (I) are shown (n = 6). (**J** to **Q**) Immune effector cells in tumors from mice treated with 200 μg of mFc or 200 μg of NDC were quantified by flow cytometry (n = 3). Scatter plots showing the proportion of cells in a specific cell population. The percentages of CD86^+^ DCs (J), MHC II^+^ DCs (K), NK cells (L), CD69^+^ NK cells (M), GB^+^ NK cells (N), CD8^+^ T cells (O), GB^+^ CD8^+^ T cells (P) and IFN-γ^+^ CD8^+^ T cells (Q) are shown based on their respective markers. (**R**) BALB/c mice were subcutaneously inoculated with 1×10^6^ of CT26 cells (induced by IFN-γ) and treated with 200 μg of mFc or 200 μg of Nb16-SZU-101 dosed with anti-murine CD4, anti-murine CD8, anti-murine NK1.1, or clodronate liposomes. Tumor weight is shown (n = 6). (**S**) Effect of 4G11 to block the inhibitory signal from RKO cells (induced by IFN-γ) to Jurkat-NFAT- luciferase cells. The data are shown in relative luciferase units (RLU). (**T** and **U**) PD-1/PD-L1 dual-humanized BALB/c mice were subcutaneously inoculated with 5×10^5^ of CT26/hPD-L1 cells. Mice bearing tumors were treated with 130 μg of hIgG1, 200 μg of 4G11 or 200 μg of 4G11-SZU-101. Tumor growth (T) in the mice and the weight of tumors (U) are shown (n = 6). The data are presented as the mean ± SEM. * p < 0.05; ** p < 0.01; *** p < 0.001; ns, not significant as determined by one-way ANOVA with multiple comparison tests or unpaired t test.

### In vivo antitumor activity of the PD-L1 and TLR7 dual-targeting NDC

SZU-101 enhanced the immunogenicity of tumors and promoted PD-L1 expression. Therefore, the PD-L1 and TLR7 dual-targeting NDC may have good inhibitory effects on tumors with different levels of PD-L1 expression and both “hot” and “cold” tumors. CT26 tumors (with or without IFN-γ induction for triggering expressing PD-L1) were treated with Nb16-SZU-101. In both CT26 tumor models, Nb16-SZU-101 significantly inhibited tumor growth, with 78.9% inhibition in the uninduced CT26 model and 83.6% in the postinduced CT26 model (Fig. 7, D-G and fig. S8, K-L). The NDC also exerted potent antitumor effects in a B16-F10 xenograft tumor model with low PD-L1 expression levels (Fig. 7, H and I and fig. S8, M-N).

### The PD-L1 and TLR7 dual-targeting NDC exerts antitumor effects by combining innate and adaptive immunity

The NDC we developed can respond to both “hot” and “cold” tumors to exert potent antitumor activity. To further illustrate the effect of the NDC, immunophenotyping was used to show that NDC treatment reshaped the tumor immune microenvironment. In CT26 tumor-bearing mice, NDC promoted the maturation of DCs compared to the isotype controls (Fig. 7, J and K). Consistent with the results of RNA-seq analysis, the NDC significantly increased the function of tumor-infiltrating cytotoxic cells, promoting their expression of granzyme B and IFN-γ (Fig. 7, L-Q). In addition, NDC administration repolarized TAMs, reduced TGF-β^+^ macrophages and increased PD-L1 expression in macrophages (fig. S9, A-D). IFN-γ^+^ CD4^+^ T cell infiltration was also significantly increased in the NDC treatment group (fig. S9E). Thus, the PD-L1 and TLR7 dual-targeting NDC plays an antitumor effect by combining innate and adaptive immunity.

Using a series of antibody-based immune cell depletion studies in CT26 tumor-bearing mice (fig. S10), we found that the antitumor efficacy of NDC was abrogated by the depletion of CD8^+^ T cells and NK cells, while the depletion of CD4^+^ T cells and macrophages did not markedly hinder the antitumor efficacy of NDC (Fig. 7R and fig. S11). Therefore, the potent antitumor effects NDC exerted relied mainly on CD8^+^ T cells and NK cells, which is consistent with our existing results. The deletion of macrophages did not significantly affect the antitumor efficacy of NDC, probably because macrophage subtypes in the tumor microenvironment are complex, with both antitumor M1 macrophages and tumor-promoting M2 macrophages. Therefore, the deletion of all macrophages is not sufficient to account for their role in the efficacy of NDC.

### The PD-L1 and TLR7 dual-targeting NDC has clinical development prospects

The experimental results of Nb16-SZU-101 treatment represent our proof-of-concept success. To make this novel drug molecule ready for clinical application, we obtained the nanobody 4G11, which targets human PD-L1 with high blocking activity (Fig. 7S). Based on the PD-L1 nanobody 4G11, we developed another PD-L1 and TLR7 dual-targeting NDC named 4G11-SZU-101 using the same method as used to prepare the Nb16-SZU-101.

Considering that both host-expressed PD-L1 and tumor cell-expressed PD-L1 may contribute to the efficacy of PD-L1 nanobodies, we chose PD-1/PD-L1 dual-humanized mice rather than PD-1 humanized mice to evaluate the in vivo antitumor activity of 4G11-SZU-101. 4G11-SZU-101 showed potent antitumor effects in a CT26/hPD-L1 xenograft tumor model established with PD-1/PD-L1 dual-humanized BALB/c mice, showing a tumor inhibition rate of 68.7% (Fig. 7, T and U and fig. S12). 4G11-SZU-101 demonstrated superior antitumor activity than single agents. Moreover, no pathological damage to the major organs of the mice was observed after 4G11-SZU-101 treatment, and no abnormalities in hematology or plasma chemistry parameters were observed, demonstrating the good safety of this NDC (Data file S1, S2 and S3). All the results demonstrated the clinical development potential of this novel drug molecule.

## DISCUSSION

The landscape for clinical trials of ICB therapies is now moving toward more efficacious combination strategies (13). Other checkpoints, costimulation molecules, antiangiogenic, epigenetic reprogramming, and other combination therapeutic strategies are being extensively studied in preclinical and clinical trials (*52*). Determinants of the response rate to PD-1/PD-L1 antibody therapy include the tumor tissue immune infiltration level, tumor mutational burden, intratumoral PD-L1 expression level, activity of antigen presentation machinery in tumors, etc. (*14, 52, 53*). Our strategy is to improve the response rate of PD-L1 antibodies by combining TLR7 agonists.

Advances in drug delivery techniques allow new drugs to be formulated that improve pharmacokinetics, enhance accumulation in solid tumors and minimize toxic effects (*54*). The molecular weight of nanobodies with better tumor penetration is one-tenth that of monoclonal antibodies (*35, 39*). To improve the in vivo activity and half-life of nanobodies, it is often necessary to attach the nanobodies to Fc fragments (*55*). Notably, Nb16, which we developed and used in this study, is only one-half the size of conventional monoclonal antibodies after the Fc fragment is attached and can still specifically accumulate in tumors with greater tumor penetration than the general PD-L1 monoclonal antibodies. Thus, nanobodies modified with Fc fragments (similar to heavy-chain-only antibodies) still have superior tumor penetration and drug delivery capabilities compared to monoclonal antibodies. Systematic administration of SZU-101 did not achieve good solid tumor targeting, and intratumoral administration is an unrealistic clinical route of administration. Although SZU-101 combined with Nb16 treatment achieved good therapeutic effects, dual-targeting NDC Nb16-SZU-101 had better antitumor activity, probably because of the precise delivery of SZU-101 to tumor tissues as facilitated by the Nb16.

PD-L1 levels in tumor tissues are generally considered to be among the major predictive markers of PD-L1 antibody therapeutic efficacy (*56*). For tumors with low levels of PD-L1, induction of PD-L1 expression may be necessary to improve the PD-L1 antibody therapeutic efficacy and tumor targeting. The TLR7 agonist SZU-101 can play this role. SZU-101 can activate innate immune cells and promote the secretion of the antitumor cytokines TNF-α and IFN-γ. We found that SZU-101 cannot directly induce PD-L1 expression in CT26 cells, except for CT26 cells cocultured with splenocytes, suggesting that the role of SZU-101 in promoting the cellular expression of PD-L1 is mainly dependent on potent PD-L1 inducers, TNF-α and IFN-γ, which are secreted by immune cells after activation. Interestingly, in vivo, SZU-101 induced higher PD-L1 expression in APCs than in tumor cells, suggesting that the NDCs we developed prefer to target host APCs for action. However, SZU-101 treatment alone may exacerbate the immune tolerance environment of the tumor. Accordingly, combining SZU-101 with the PD-L1 nanobody for treatment can overcome the effects of an enhanced tumor immunosuppressive environment associated with SZU-101 administration while also improving the efficacy and response rate of PD-L1 nanobody therapy.

In addition, although PD-L1 nanobodies can attenuate the immunosuppression of T cells, the absence of a sufficient innate immune response may be a limiting factor that restricts the development of an effective adaptive antitumor immune response, thus triggering the formation of “cold” tumors (*52*). Currently, the development of innate immune agonists is a hot research topic, and drugs targeting stimulator of interferon genes (STING), cyclic GMP-AMP synthase (cGAS) and TLRs are showing good progress in tumor therapy (*52*). The TLR7 agonist SZU-101 that we developed can activate innate immune cells such as dendritic cells and NK cells and induce the upregulation of MHC II expression by APCs and upregulation of MHC I expression by tumor cells, thus achieving adequate activation of the innate immune system, promoting the activity of intratumoral antigen-presenting machinery, increasing the immunogenicity of tumors, achieving more effective adaptive immunosensitization, and transforming “cold” tumors into “hot” tumors. Moreover, SZU-101 can induce high secretion of the antitumor cytokines TNF-α and IFN-γ from immune cells in vivo and promote macrophage repolarization to improve the tumor immune microenvironment.

To the best of our knowledge, the Nb16-SZU-101 we developed is the first PD-L1 and TLR7 dual-targeting NDC reported to date. The potent antitumor efficacy of Nb16-SZU-101 also supports our concept of combining a PD-L1 antibody and TLR7 agonist to attenuate tumor immune tolerance and promote the activity of antigen-presenting machinery, which can lead to the activation of both innate and adaptive immunity to achieve the desired antitumor effect. The experimental results of 4G11-SZU-101 treatment demonstrated the clinical development potential of this novel drug molecule.

Of course, our study has many limitations, and further research is needed. First, the nanobodies Nb16 and 4G11 did not show potent antitumor effects, which may be related to the affinity and action epitopes of these nanobodies. Subsequent optimization, such as affinity maturation modification of nanobodies, is needed. Second, the NDC we developed currently functions through a nondeterministic coupling technique, and it is necessary to subsequently adopt a deterministic coupling technique and optimize the DAR value of the NDC. It is expected that by screening PD-L1 nanobodies with better antitumor activity and optimizing the appropriate DAR values, PD-L1 and TLR7 dual-targeting NDCs will achieve better antitumor effects. Third, we have evaluated the antitumor activity of NDC only on PD-1/PD-L1 dual-humanized BALB/c mice thus far, and therefore, the inhibitory effect of NDCs need to be tested on human-derived tumor growing in immune system–humanized mice. In addition, the role of NDC in tumor growth needs to be further investigated to identify targeted drugs that can be combined with NDC to achieve better antitumor efficacy.

In conclusion, our study reveals that the TLR7 agonist SZU-101 and the PD-L1 nanobody Nb16 or 4G11 can exert synergistic antitumor effects. The combination of the PD-L1 nanobody and a TLR7 agonist shows a notable rationale (Fig. 8). First, PD-L1 nanobodies have the desired tumor penetration and tumor tissue targeting abilities such that the TLR7 agonists can be precisely delivered to tumor tissues. Second, TLR7 agonists can induce high expression of PD-L1 within tumors; therefore, PD-L1 nanobodies can achieve better antitumor effects and responses to tumors with low levels of PD-L1. Third, the lack of an effective mechanism to activate innate immunity makes it difficult for PD-L1 nanobodies to fully restore T-cell antitumor immune responses. TLR7 agonists can enhance tumor immunogenicity and convert “cold” tumors into “hot” tumors by activating innate immunity and promoting the activity of antigen-presenting machinery within the tumors. Accordingly, we have developed a type of PD-L1 and TLR7 dual-targeting NDC with potent antitumor activity, for the first time, and 4G11-SZU-101 was developed with potential for clinical development. The PD-L1 and TLR7 dual-targeting NDC represents a combination therapeutic strategy that coordinates innate and adaptive immunity to elicit a potent and broadly responsive antitumor effect.

**Fig. 8.**
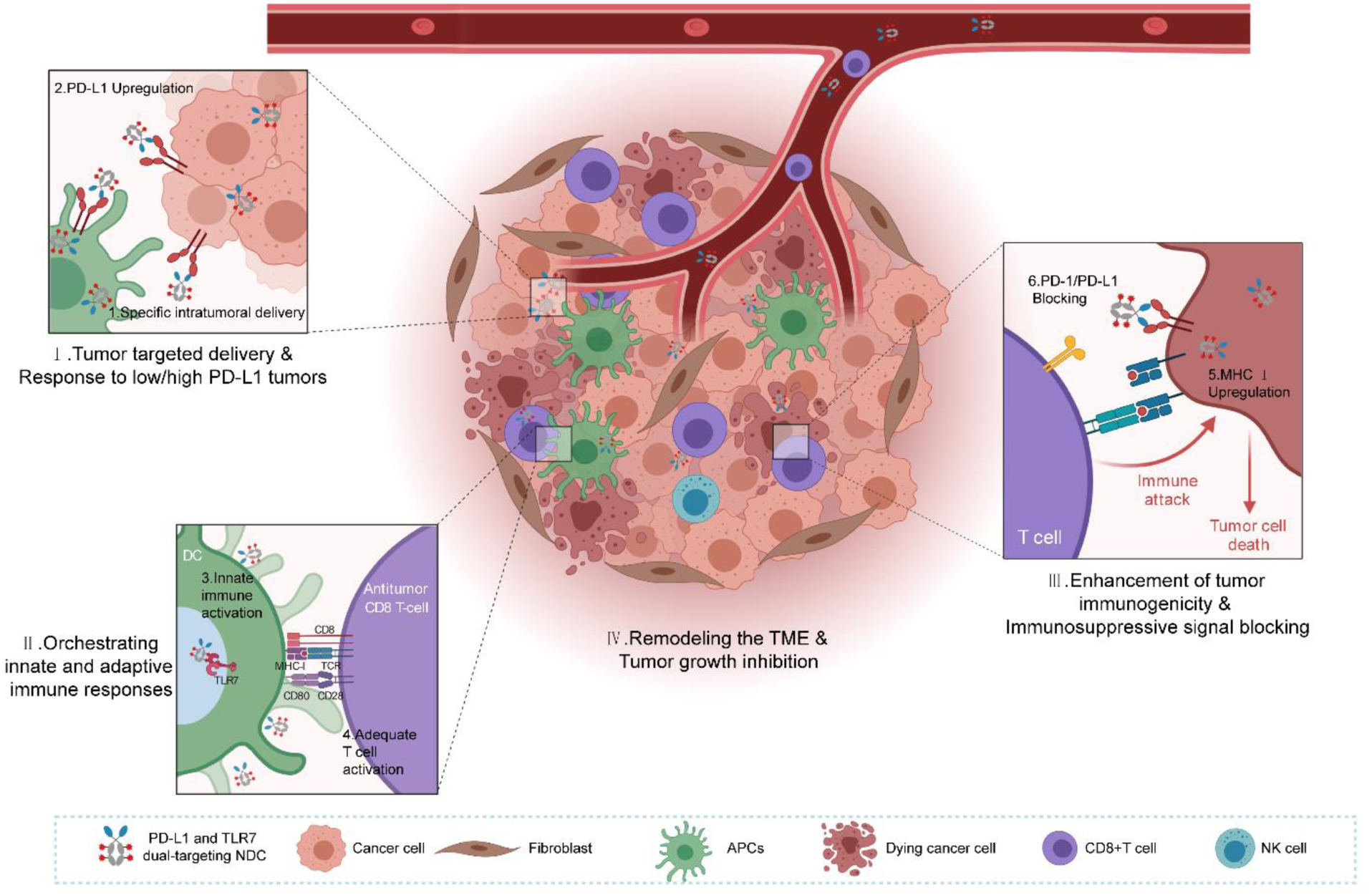
The PD-L1 and TLR7 dual-targeting NDC mechanism of action diagram. The NDC can utilize the complementary effects of PD-L1 nanobodies and TLR7 agonists to fully mobilize the innate and adaptive antitumor immune responses. PD-L1 nanobodies can deliver TLR7 agonists to tumor tissues, and TLR7 agonists can promote PD-L1 expression in tumor tissues, achieving a response to tumors with low PD-L1 expression. TLR7 agonists can activate the innate immune responses and thus assist in the full activation of adaptive immunity. TLR7 agonists can enhance the immunogenicity of tumors, and PD-L1 nanoantibodies can release the immunosuppression of T cells by tumors. The figure was created with BioRender.com.

## MATERIALS AND METHODS

### Study design

This study was designed to characterize the rationalization of PD-L1 nanobody and TLR7 agonist combination therapy and the therapeutic potential of PD-L1 and TLR7 dual-targeting NDCs. We produced and characterized nanobodies (Nb16 and 4G11), a TLR7 agonist small molecule (SZU-101) and NDCs (Nb16-SZU-101 and 4G11-SZU-101). The in vitro and ex vivo activity of the drugs was evaluated in multiple cell models. The antitumor effects, immune responses and mechanism of action were tested in several mouse models, including a PD-1/PD-L1 dual-humanized BALB/c mouse models. The animals were randomly assigned to experimental groups to ensure that the mean xenograft tumor volume and mean mouse body weight were similar across groups, but experimenters were not blinded to the groups. The number of mice per group in each experiment was determined to ensure statistical power. Six to nine animals in each group were used for the antitumor efficacy studies. The number of mice in each group and the statistical tests used for each experiment are described in each figure legend.

### Statistical analyses

Statistical analysis was performed using GraphPad Prism 8 Software. Statistical methods and sample sizes in the experiments are described in each figure. P values < 0.05 were considered to be significant. For all other Materials and Methods, see the Supplementary Materials.

## Acknowledgments

We thank H. Wang, Y. Hang, M. Zhang, and the staff of GemPharmatech for their help with mouse husbandry and experiments. We thank Professor Y. Huang for scientific discussions. We thank the staff of the Center for Drug Safety Evaluation and Research (CDSER), Shanghai Institute of Materia Medica (SIMM) for the safety evaluation. We thank Conjugenix Pharma-tech Co., Ltd. (Shenzhen, China) for assisting molecule synthesis. We also thank Professor Y.Geng for kindly providing the nanobody 4G11.

## Author contributions

L.G., G.J., X.Y. and Y.L. designed the experiments and analyzed the data. X.Y. and Y.L. performed the experiments and prepared the manuscript. B.C., Y.T., Y.C. and P.X. assisted in performing the experiments and preparation of the manuscript. X.J. expressed and purified the nanobody 4G11. W.H. and F.T. performed the identification for NDC by mass spectrometer. J.Z. performed the synthesis of SZU-101 and SZU-101-NHS. Y.W., J.R. and J.S. assisted in data interpretation.

## Data and materials availability

The BioProject accession number for the mouse RNA-seq data is PRJNA706848 (https://www.ncbi.nlm.nih.gov/sra/PRJNA706848). Other data are available in the main text or the supplementary materials. Correspondence and requests for materials should be addressed to L.G.

## MATERIALS AND METHODS

### Cell lines

FreeStyle™ HEK 293F was purchased from Thermo Fisher and grown in FreeStyle™ 293 expression medium (Thermo Fisher). HEK 293T, Jurkat, RKO, CT26 and B16-F10 cell lines were purchased from the American Type Culture Collection (ATCC). HEK 293T and B16-F10 cells were cultured in Dulbecco’s Modified Eagle Medium (DMEM; Meilunbio) supplemented with 1% penicillin/streptomycin (Invitrogen) and 10% heat-inactivated fetal bovine serum (FBS; Life Technologies). Jurkat cells were cultured in RPMI1640 medium (Meilunbio) containing with 1% penicillin/streptomycin and 10% FBS. CT26 cells were cultured in RPMI1640 medium containing with 1% penicillin/streptomycin, 10% FBS, 1mM sodium pyruvate (Invitrogen) and 4.5 g/L glucose (Invitrogen). RKO cells were cultured in Minimum Essential Medium (MEM; Meilunbio) supplemented with 1% penicillin/streptomycin and 10% FBS. NK-92 cell lines were purchased from Procell and cultured with special medium for NK-92 cells (Procell). CT26/hPD-L1 cell lines were provided by GemPharmatech. All cells were cultured at 37°C in a 5% CO2 humidified atmosphere.

### Plasmids

Vector pFUSE-hIgG1-Fc2 and pFUSE-mIgG2b-Fc were purchased from Invivogen. Vector pLVX-puro was purchased from Clontech. Vector pLV-NFAT-RE-Luci was purchased from Inovogen. pVSV-G, pMDlg/pRRE, and pRSV-Rev were purchased from Youbio. pECMV-PD-1/PDCD1 was was purchased from Miaolingbio.

### Mice

Female (Six- to eight-week-old) BALB/c and C57BL/6J mice were purchased from Shanghai Sipp-BK laboratory animal Co. Ltd. All mice were maintained under specific pathogen-free (SPF) conditions in the animal facility of the Shanghai Institute of Materia Medica, Chinese Academy of Sciences. BALB/c-hPD1/hPDL1 mice were purchased from GemPharmatech Co., Ltd and housed in SPF condition (GemPharmatech). Animal care and experiments were performed in accordance with GemPharmatech and the Shanghai Institute of Materia Medica, Chinese Academy of Sciences, using protocols approved by the Institutional Laboratory Animal Care and Use Committee (IACUC).

### Expression and purification of murine PD-L1 and human PD-L1

Genes encoding mouse PD-L1 (mouse PD-L1, amino acid residues 19-239) and human PD-L1 (amino acid residues 19-238) were cloned into pFUSE-hIgG1-Fc2 expression vector (Invivogen). The recombinant plasmids were used for transfection and PD-L1 expression. Murine PD-L1-Fc and human PD-L1-Fc were expressed by transfecting FreeStyle™ HEK 293F cells. Briefly, FreeStyle™ HEK 293F cells were cultured in serum-free FreeStyle™ 293 expression medium in an orbital shaker incubator at 37°C and 120 rpm with 8% CO_2_. FreeStyle™ HEK 293F cells (1.0 × 10^6^ cells/mL) were transfected with a mixed solution [plasmids, PEI (Polyscience) and PBS], which has been incubated at room temperature for at least 20 min. Transfected FreeStyle™ HEK 293F cells were allowed to secrete proteins for four to five days. Then the medium was collected for PD-L1 purification using immobilized protein A beads (Bestchrom). The purity and size of the PD-L1 were identified using 15% sodium dodecyl sulfate polyacrylamide gel electrophoresis (SDS-PAGE), followed by staining with coomassie brilliant blue.

### Acquisition of PD-L1 nanobodies

The nanobodies Nb16 and 4G11 were obtained in the same way as in our previous study (33), including camel immunization, nanobody library construction and screening of nanobodies by phage display technology.

### Generation and purification of nanobodies in FreeStyle™ HEK 293F Cells

The VHHs of Nb16 and 4G11 were inserted into pFUSE-mIgG2b-Fc or pFUSE-hIgG1-Fc2 vector respectively. The plasmids of VHH-pFUSE-mIgG2b-Fc and VHH-pFUSE-hIgG1-Fc2 were transfected into FreeStyle™ HEK 293F cells to express the nanobodies. The transfected FreeStyle™ HEK 293F cells were cultured for protein expression in the FreeStyle™ 293 expression medium for four to five days. Then the medium was collected for nanobodies purification using immobilized protein A beads. The SDS-PAGE was used to characterize the nanobodies (15 kDa) with an Fc tag (27 kDa).

### Antibody conjugation and characterization

The synthesis of SZU-101 was performed as in our previous study (25). SZU-101, 1-ethyl-3(3-dimethylpropylamine) carbodiimide (EDCI) and N-hydroxysuccinimide (NHS) were dissolved in dimethyl sulfoxide (DMSO), then stirred at room temperature for three hours. The reaction process was monitored by liquid chromatography-mass spectrometry (LC-MS, Waters). After the reaction was finished, ten times the amount of double-distilled water was added. After filtered, the products were dried under vacuum to obtain SZU-101-NHS activated ester.

Subsequently, the activated ester was dissolved with DMSO. Antibodies and small molecules were reacted in a molar ratio of 1:10. Small molecules were added to Nb16 or 4G11 solution and the reaction was stirred at 4°C for 4 hours. After the reaction, PBS was added to the mixture. After mixed, the small molecules were removed by filtration with a 10 kD biofilter membrane to obtain the novel compound Nb16-SZU-101 or 4G11-SZU-101. To identify NDC, the antibodies were denatured to open the disulfide bond, after which the sample was identified by Xevo G2-XS QTOF mass spectrometer (Waters). The DAR value was calculated mainly based on the increase in molecular weight of NDC over the uncoupled antibody.

### Generation of murine PD-L1 stably expressed HEK 293T cells

Stable expression cell lines can be achieved by using lentiviral vector. Briefly, murine PD-L1 full gene was subcloned into pLVX-puro lentiviral expression vector (Clontech), which was then transfected into HEK 293T cells with the package plasmids. Then the virosomes were collected and inflected into HEK 293T cells. The murine PD-L1 stably expressed HEK 293T cells (293T/ mPD-L1 cells) were obtained after puromycin resistance selection. To check whether the stable cell lines were constructed, 293T/ mPD-L1 cells (3×10^5^ cells) were incubated with the specific anti mPD-L1 Nb-human Fc proteins (1 μg). Goat anti-Human IgG Fc (FITC) antibody (Abcam) was acted as secondary antibody. Data were obtained by flow cytometry (BD Biosciences).

### Preparation of mouse PD-1-biotin

Mouse PD-1 protein (1 mg/mL) and biotin (10 mM) were reacted at 4°C for 2 h. The solution was mixed three times during the reaction. After the reaction, the solution was ultrafiltered using an ultrafiltration tube and the concentration was adjusted to 1 mg/mL.

### Determination of PD-L1 binding affinity and PD-1/PD-L1 blocking activity of nanobodies and NDC

#### Measurement of murine PD-L1 binding affinity of Nb16

EC50 value of Nb16 was measured by standard ELISA methods. 96-well ELISA plates (Nunc) were coated with mouse PD-L1-Fc (2 μg/mL) in PBS overnight at 4°C. After being washed five times with PBST buffer (PBS with 0.05% Tween-20), 96- well plates were blocked with 200 μL of 1% BSA solution for two hours at room temperature. After being washed, 96-well plates were added with 100 μL of serial dilutions of Nb16 (starting at 50 μg/mL) and incubated for one hour at room temperature. After being washed, 96-well plates were incubated with 100 μL of anti-mFc-horseradish peroxidase (HRP, Thermo Fisher) for one hour at room temperature. After being washed, 96-well plates were incubated with 100 μL of tetramethyl benzidine (TMB) substrate solution (Thermo Fisher) at room temperature for 10 min. The reaction was terminated by adding 2 M of sulfuric acid solution, and the absorbance value was measured at 450 nm with an automatic microplate reader SpectraMax (Molecular Devices).

#### Measurement of murine PD-1/PD-L1 blocking activity of Nb16

IC50 value of Nb16 was measured by standard competition ELISA methods. 96-well ELISA plates were coated with mouse PD-L1-Fc (2 μg/mL) in PBS overnight at 4°C. After being washed and blocked, 96-well plates were added with 50 μL of serial dilutions of Nb16 (starting at 50 μg/mL) and mouse PD-1-biotin (50 μL, 10 μg/mL) and incubated for one hour at room temperature. After being washed, 96-well plates were incubated with 100 μL of anti-streptomavidin-HRP (Abcam) for one hour at room temperature. After being washed, 96-well plates were incubated with 100 μL of TMB substrate solution at room temperature for 10 min. The reaction was terminated by adding 2 M of sulfuric acid solution, and the absorbance value was measured at 450 nm.

#### Detection of Nb16 binding to murine PD-L1 stably expressed HEK 293T cells

100 μL of 293T/ mPD-L1 cells (2.5 × 10^5^) and 2 μg/mL of Nb16 were incubated at 4°C for 20 min. After being washed, cells were stained with 3 μg/mL of goat anti-human IgG-Fc (FITC, Thermo Fisher) solution for 20 min at 4°C. After being washed, data were obtained by flow cytometry (BD Biosciences).

#### Detection of PD-1/PD-L1blocking activity of Nb16 on murine PD-L1 stably expressed HEK 293T cells

100 μL of 293T/ mPD-L1 cells (2.5 × 10^5^), 2 μg/mL of Nb16 and 10 μg/mL mouse PD-1-biotin were incubated at 4°C for 20 min. After being washed, cells were stained with 3 μg/mL of goat anti-streptomavidin PE (Thermo Fisher) solution for 20 min at 4°C. After being washed, data were obtained by flow cytometry.

#### Measurement of human PD-L1 binding affinity of 4G11

The binding kinetics of nanobody 4G11 for human PD-L1 was analyzed using Octet-Red96 (Pall ForteBio) biolayer interferometry. Purified nanobodies 4G11 were immobilized on activated protein A biosensor at a concentration of 20 μg/mL and tested with a two-fold gradient dilution of human PD-L1 into the sample, and response values were recorded. Fitted curves were constructed with ForteBio data analysis software.

#### Measurement of human PD-1/PD-L1 blocking activity of 4G11

Using surface plasmon resonance technique, human PD-1 protein was coupled to CM5 chip by amino carboxyl coupling and incubated with high concentration of nanobody and human PD-L1. 100 nM and 400 nM of human PD-L1 were selected, and the concentrations of 4G11 were 400 nM, 200 nM, 100 nM …0 nM in order. The samples were incubated at room temperature for 30 min and the incubated samples were injected with Biacore T200 (GE) for measurement.

#### Measurement of mouse PD-L1 binding affinity of Nb16-SZU-101

100 μL of 293T/ mPD-L1 cells (2.5 × 10^5^) and serial dilutions of Nb16-SZU-101 (100, 50, 25, 12.5, 6.25, 3.12, 1.56, 0.78, 0.39, 0.20, 0.10, 0.05, 0.02, 0.01, 0.006, 0.003 and 0.0008 μg/mL) were incubated at 4°C for 20 min. After being washed, cells were stained with 3 μg/mL of goat anti-human IgG-Fc (FITC) solution for 20 min at 4°C. After being washed, data were obtained by flow cytometry.

#### Measurement of human PD-1/PD-L1 blocking activity of Nb16-SZU-101

100 μL of 293T/ mPD-L1 cells (2.5 × 10^5^), serial dilutions of Nb16-SZU-101 (100, 50, 25, 12.5, 6.25, 3.12, 1.56, 0.78, 0.39, 0.20, 0.10, 0.05, 0.02, 0.01, 0.006, 0.003 and 0.0008 μg/mL) and 20 μg/mL of mouse PD-1-biotin were incubated at 4°C for 20 min. After being washed, cells were stained with 3 μg/mL of goat anti-streptomavidin (PE) solution for 20 min at 4°C. After being washed, data were obtained by flow cytometry.

### In vivo treatment study

#### Evaluation of the antitumor activity of Nb16

CT26 cells were cultured and had passed at least 3 generations before tumor bearing, and CT26 cells were induced with mouse IFN-γ (GenScript) at a final concentration of 100 ng/mL for 24 h, resulting in high expression of PD-L1 on the cell surface. CT26 tumor cells (1×10^6^ cells) were subcutaneously inoculated into each female BALB/c mouse. Mice were randomly grouped. The mFc (200 μg) and Nb16 (200 μg) were administered intraperitoneally on days 1, 5, 9, 12 and 14. Tumors were measured on days 4, 9, 12, 13 and 14, and tumor volume was calculated as tumor volume (mm^3^) = tumor length × width × width / 2, and tumor growth curves were plotted. Mice were sacrificed on day 15.

#### Evaluation of the antitumor activity of SZU-101

CT26 cells were cultured and had passed at least 3 generations at the time of tumor bearing. CT26 tumor cells (1×10^6^ cells) were subcutaneously inoculated into each female BALB/c mouse. Mice were randomly grouped. SZU-101 (3mg/kg) was administered intraperitoneally or intratumorally every two days from day 2, for a total of 8 times. Tumors were measured as mentioned above. Mice were sacrificed on day 16 and the tumor tissues were obtained.

#### Evaluation of the antitumor activity of Nb16 and SZU-101 combination treatment

CT26 cells were cultured and had passed at least 3 generations at the time of tumor bearing, and CT26 cells were induced with mouse IFN-γ (100 ng/mL) for 24 h. CT26 tumor cells (1×10^6^ cells) were subcutaneously inoculated into each female BALB/c mouse. Mice were randomly grouped into 4 groups of 8 mice each, including the mFc group, the Nb16 group, the SZU-101 group, and the combination group. Nb16 was administered intraperitoneally at a dose of 200 μg/each on days 1, 5, 9, 12 and 14 for a total of 5 doses. SZU-101 was administered peritumorally at a dose of 3 mg/kg/each from day 9 to day 14. Tumors were measured on days 4, 9, 12, 13 and 14. Mice were sacrificed on day 15 and the tumor tissues were obtained.

#### Evaluation of the antitumor activity of Nb16-SZU-101 for CT26 (IFN-γ induced) model

CT26 cells were cultured and had passed at least 3 generations at the time of tumor bearing, and CT26 cells were induced with mouse IFN-γ (100 ng/mL) for 24 h. CT26 tumor cells (1×10^6^ cells) were subcutaneously inoculated into each female BALB/c mouse. Mice were randomly divided into four groups of six mice each, including the mFc group, the Nb16 group, the combination group and the Nb16-SZU-101 group. Dosed on days 2, 6, 10 and 13 for a total of 4 doses, mFc, Nb16 and Nb16-SZU-101 were administered intraperitoneally at a dose of 10mg/kg/each, SZU-101 was administered intraperitoneally at a dose of 0.5 mg/kg/each. Tumors were measured on days 7, 9, 10, 11, 12, 13 and 14. Mice were sacrificed on day 14 and the tumor tissues were obtained.

#### Evaluation of the antitumor activity of Nb16-SZU-101 for CT26 (uninduced) model

Six- to eight-week-old BALB/c mice were randomly divided into 2 groups. Mice were inoculated subcutaneously with CT26 cells (1×10^6^). The mFc and Nb16-SZU-101 were administered intraperitoneally with 10 mg/kg/each, respectively. Each group was administered four times on days 2, 5, 8 and 11 after tumor bearing. Tumors were measured on days 7, 8, 9, 10, 11 and 12. Mice were euthanized on day 12 and the tumor tissues were obtained.

#### Evaluation of the antitumor activity of Nb16-SZU-101 for B16-F10 model

Six- to eight-week-old C57BL/6 mice were randomly divided into 2 groups. Mice were inoculated subcutaneously with B16-F10 cells (1×10^6^). The mFc (10 mg/kg) and Nb16-SZU-101 (10 mg/kg) were administered intraperitoneally. Each group was administered four times on days 2, 5, 8 and 11 after tumor bearing. Tumors were measured on days 7, 8, 9, 10, 11 and 12. Mice were euthanized on day 12 and the tumor tissues were obtained.

#### Evaluation of the antitumor activity of 4G11-SZU-101 for CT26/hPD-L1 xenograft tumor model in PD-1/PD-L1 dual-humanized BALB/c mice

Six- to eight-week-old dual-humanized BALB/c mice were inoculated with CT26/hPD-L1 tumor cells (5×10^5^ cells, 100 μL) subcutaneously on the right side of the back of the mice above the thigh. When the average tumor volume reached 80-120 mm^3^, the mice were randomly divided into three groups (hIgG1 group, 4G11 group and 4G11-SZU-101 group) according to the tumor volume, with six mice in each group. The day of grouping was defined as day 0, and dosing was started on day 0. hIgG1 was administered at a dose of 130 μg/each, while 4G11 and 4G11-SZU-101 were administered at a dose of 200 μg/each. The dose was performed every three days for a total of six doses, during which tumor growth was observed and recorded. After the final administration, the mice were euthanized. Tumor volume was calculated as tumor volume (mm^3^) = tumor length × width × width / 2, and tumor growth curves were plotted.

### In vivo imaging

CT26 tumor-bearing mice were injected intraperitoneally with fluorescein Cy7- labeled Nb16 and fluorescein Cy7-labeled anti-mouse PD-L1 monoclonal antibody 10F.9G2. Fluorescent antibody distribution in vivo was monitored using an IVIS imaging system (Caliper PerkinElmer) at predetermined time points (0 h, 1 h, 2 h, 4 h and 7 h). At the end of the experiment, mice were sacrificed, tumors were obtained and imaged in vitro using the IVIS imaging system.

### TCGA cohorts analysis

The Tumor Immune Estimation Resource (TIMER2.0, http://timer.comp-genomics.org/) online tool was used to analyze TLR7 expression levels in the Cancer Genome Atlas (TCGA) datasets. And TIMER2.0 was used to explore the prognostic value of TLR7 in patients with different types of cancers, while the data were showed as Kaplan-Meier curves. We also used TIMER2.0 to analyze the correlation between TLR7 expression levels in TCGA datasets and the intratumoral infiltration levels of DCs, macrophages, neutrophils, tumor-associated macrophages, CD4^+^ T cells and CD8^+^ T cells. In addition, we also analyzed the association between TLR7 expression levels and TNF, IFNG, CD274, HLA-DRA and HLA-DRB1 in multiple tumor datasets of TCGA by TIMER2.0. Heat maps were drawn by GraphPad Prism 8 Software.

### Jurkat-NFAT effector cells reporter assay

Jurkat-NFAT-luciferase cells were constructed by stable transfection of pLV-NFAT-RE-Luci by lentivirus. Jurkat-NFAT-luciferase cells were stimulated with 50 ng/mL of phorbol-12-myristate-13-acetate (PMA) and 1 µg/mL of ionomycin for 18 h.

To test the blocking activity of Nb16, plasmids pECMV-PD-1/PDCD1 were transfected into Jurkat-NFAT-luciferase cells by PEI. In a 24-well plate (Corning), each well was added with CT26 cells (300μL, 3×10^5^ cells/mL; 100 ng/mL mouse IFN-γ induced for 24 h). After 24 h, Jurkat-NFAT-luciferase cells (300μL, 3×10^5^ cells/mL) were added into the wells.

To test the blocking activity of 4G11, in a 24-well plate, each well was added with RKO cells (300μL, 3×10^5^ cells/mL; 100 ng/mL human IFN-γ induced for 24 h). After 24 h, Jurkat-NFAT-luciferase cells (300μL, 3×10^5^ cells/mL) were added into the wells.

After co-culture of cells for 24 h, cells were collected and 120 μL of cell lysate (Yeasen) was added. The lysate after sufficient lysis was centrifuged at 10,000-15,000 rpm for three to five min. 100 μL of firefly luciferase assay reagent (Yeasen) was added to the supernatant. The solution was mixed 2-3 times (no vortex mixing) before measuring the RLU (relative light unit) in an automatic microplate reader SpectraMax (Molecular Devices).

### In vitro cell assays

#### SZU-101 activated NK cells

In a 48-well plate (Thermo Fisher), each well was added with NK-92 cells (300 μL, 5×10^5^ cells/mL) and added with 50 μM of SZU-101. After 24 h, the NK-92 cells were collected and blocked with 4% FBS and anti-CD16/CD32 (FcγRIII/FcγRII, 2.4G2, BD Biosciences) for 1 h at 4°C. Then the NK-92 cells were stained with anti- human CD25 APC (BD Biosciences) and anti-human CD69 APC-Cy7 (BD Biosciences) for 20 min at 4°C. CD25 and CD69 signals of NK-92 cells were detected by flow cytometry (ACEA NovoCyte).

#### SZU-101 did not promote PD-L1 expression in CT26 cells

In a 48-well plate, each well was added with CT26 cells (300 μL, 3×10^5^ cells/mL) and added with 50 μM of SZU-101. After 24 h, the CT26 cells were collected and blocked with 4% FBS and anti-CD16/CD32 (FcγRIII/FcγRII, 2.4G2) for 1 h at 4°C. Then the CT26 cells were stained with Ms CD274 BV650 MIH5 (BD Biosciences) for 20 min at 4°C. PD-L1 signals of CT26 cells were detected by flow cytometry (ACEA NovoCyte).

### Ex vivo analysis of immune responses

#### SZU-101 promoted proliferation and cytokine secretion of splenocytes

After euthanasia, BALB/c mice were soaked in 75% alcohol for 5 min and its spleen was taken out under aseptic conditions. Single-cell suspensions of splenocytes were obtained by grinding spleen tissues in RPMI1640 medium with a syringe rubber pad, incubating with erythrocyte lysis buffer (Yeasen), and then filtering through a 70-μm nylon cell filter. Splenocytes (500 μL, 1×10^6^ cells/mL) were added to 24-well plates. Mouse splenocytes were stimulated with SZU-101(1 μM, 5 μM, 10 μM, 50 μM or 100 μM) for 24 h. The proliferation level of splenocytes was detected by Cell Proliferation Kit I (MTT, Roche). Mouse splenocytes were stimulated with SZU-101 (5 μM or 50 μM) for 24 h. TNF-α and IFN-γ secreted by splenocytes were detected by the corresponding ELISA kits (Excell Bio).

#### SZU-101 acted on bone marrow-derived DCs

After euthanasia, BALB/c mice were soaked in 75% alcohol for 5 min. Femur and tibia was taken out under aseptic conditions. Bone marrow was obtained by repeated flushing from the femur and tibia and incubating with erythrocyte lysis buffer, and then filtering through a 70-μm nylon cell filter. Bone marrow cells were cultured with DMEM medium containing 25 ng/mL of GM-CSF (GenScript) and 10 ng/mL of IL-4 (GenScript) for 8 days. Subsequently, immature DC cells were collected and stimulated with 1 μg/mL of lipopolysaccharide (LPS) or 50 μM of SZU-101 for 24 h. Maturation markers in DC cells were identified by flow cytometry. The fluorescent antibodies used included PE Hamster Anti-Mouse CD80 (BD Biosciences), APC Rat anti-Mouse CD86 (BD Biosciences) and BB515 Rat Anti-Mouse I-A/I-E (BD Biosciences).

#### SZU-101 acted on bone marrow-derived macrophages

Bone marrow cells were obtained as described above. Bone marrow cells were cultured with DMEM medium containing 20 ng/mL of GM-CSF for 4 days. The above cells were induced with 20 ng/mL of IFN-γ (GenScript) and 100 ng/mL of LPS and induced to differentiate into M1-type macrophages after one day, or the above cells were induced with 40 ng/mL of IL-4 and induced to differentiate into M2-type macrophages after one day. M2-type macrophages were stimulated with 50 μM of SZU-101 for 24h. M1 and M2 markers in macrophages were identified by flow cytometry. The fluorescent antibodies used included PE-Cyanine7 Rat Anti-Mouse F4/80 (eBioscience), APC Rat anti-Mouse CD86 (BD Biosciences) and FITC anti-mouse CD206 (BD Biosciences).

#### SZU-101 acted on spleen-derived DCs

Splenocytes were obtained as described above. 1×10^6^ cells/mL of splenocytes were added to 24-well plates as 500 μL/well. Mouse splenocytes were stimulated with SZU-101 (50 μM) for 24 h. PD-L1 levels in DCs were identified by flow cytometry. The fluorescent antibodies used included BV786 Hamster Anti-Mouse CD11c (BD Biosciences), BV510 Rat Anti-Mouse I-A/I-E (BD Biosciences) and Ms CD274 BV650 MIH5 (BD Biosciences).

#### SZU-101 acted on spleen-derived macrophages

Splenocytes were obtained as described above. 1×10^6^ cells/mL of splenocytes were added to 24-well plates as 500 μL/well. Mouse splenocytes were stimulated with 50 μM of SZU-101 for 24 h. PD-L1 levels in macrophages were identified by flow cytometry. The fluorescent antibodies used included BV421 Rat Anti-CD11b (BD Biosciences), PE Rat Anti-Mouse F4/80 (BD Biosciences) and Ms CD274 BV650 MIH5 (BD Biosciences).

#### SZU-101 acted on CT26 cells and splenocytes co-culture system

Splenocytes were obtained as described above. Splenocytes (300 μL, 5×10^5^ cells/mL) and CT26 cells (300 μL, 5×10^5^ cells/mL) were added into 24-well plates to co-culture. Cells were stimulated with 50 μM of SZU-101 for 24 h. PD-L1 levels in CT26 cells were identified by flow cytometry. The fluorescent antibodies used included Ms CD45.2 APC-Cy7 (BD Biosciences) and Ms CD274 BV650 MIH5 (BD Biosciences).

### Immune cell depletion study

Six- to eight-week-old BALB/c mice were randomly divided into 6 groups, with 6 animals in each group. Mice were inoculated subcutaneously with CT26 cells (1×10^6^). The corresponding groups were mFC group, Nb16-SZU-101 group, Nb16-SZU-101+ anti-CD4 group, Nb16-SZU-101+ anti-CD8 group, Nb16-SZU-101+ anti-NK1.1 group, and Nb16-SZU-101+ chlorophospholiposome group. Immune cells were deleted by intraperitoneal injection of antibody drugs [anti-CD4 (Biolegend, 100457), anti-CD8 (Biolegend, 100763), anti-NK1.1 (Biolegend, 108759)] on day 3 (200 μg/each) / day 4 (100 μg/ each) / day 5 (100 μg/ each) to delete the corresponding immune cells (CD4^+^ T cells, CD8^+^ T cells, NK cells), respectively, or intraperitoneal injection of chlorophosphate-liposomes (Yeasen) to remove macrophages. Immune cell deletion effect identified by flow cytometry. On day14, mice were euthanized and the tumor tissues were obtained.

### Immunophenotype analysis of tumor-bearing mice

To study the effect of SZU-101 on CT26 tumors, mice were sacrificed and tumors were harvested on day 16. To quantify the immune profiling of CT26 tumors in combination therapy study, mice were sacrificed and tumors were harvested on the end of treatment. Specifically, Nb16 was administered intraperitoneally at a dose of 200 μg/each on day 12, 14, 16, and 19 for a total of 4 doses, SZU-101 was administered peritumorally at a dose of 3 mg/kg/each from day 13 to day 19. The animals were dissected on Day 20. To quantify the immune profiling of CT26 tumors in Nb16-SZU-101 therapy study, mice were sacrificed and tumors were harvested on day 12.

For the preparing of tumor cell suspensions, subcutaneous tumor tissues were harvested and finely minced with sterile scissors. 150 mg of tumor tissues was cut into mush, digested in hyaluronidase (Yeasen) and collagenase IV (Yeasen) for 1.5 h at 37°C with 180 rpm shaking, and processed into single cell suspensions. The pretreated cell suspension was divided into two portions, one for immunophenotyping of macrophages and dendritic cells and one for immunophenotyping of lymphocytes.

Macrophage and dendritic cell samples were treated with 2-4 mL of sterile erythrocyte lysis solution (Yeasen) for 8 min and terminated with PBS buffer to remove erythrocytes and debris from the sample suspension. Lymphocyte samples were processed by centrifugation with lymphocyte separation solution (Dakewei) to obtain the lymphocyte separation layer.

Samples were blocked with 4% FBS and anti-CD16/CD32 (FcγRIII/FcγRII, 2.4G2), incubated with surface marker antibodies for 20 minutes at 4°C and then permeabilized with BD Cytofix/Cytoperm buffer before intracellular labeling antibodies were added for 30 minutes at 4°C. Then transcription factors were labeled for 45 minutes at 4°C by antibodies after permeabilized with BD TF Fix/Perm buffer. Flow cytometry analysis was performed using BD FACS Calibur or ACEA NovoCyte. Cellular events were first gated by forward and side scatter characteristics and then by characteristic fluorescence. Data processing was done through Flowjo software or NovoExpress software.

Antibody staining of tumor tissues cells for flow cytometry analysis was performed following the antibody manufacturer’s recommendations. The antibodies used in immunotyping include FITC Rat Anti-Mouse CD45 (BD Biosciences), PE-Cyanine7 Rat Anti-Mouse F4/80 (BD Biosciences), APC Rat anti-Mouse CD86 (BD Biosciences), PE Rat Anti-Mouse CD274 (BD Biosciences), Ms CD206 Alexa 647 MR5D3 (BD Biosciences), anti-mouse LAP (TW7-16B4) PE (Bioscience), FITC Hamster Anti-Mouse CD3e (BD Biosciences), PerCP-Cy™5.5 Rat Anti-Mouse CD4 (BD Biosciences), APC Rat Anti-Mouse CD8a (BD Biosciences), PE Rat Anti-Mouse IFN-γ (BD Biosciences), anti-mouse granzyme B (NGZB) PE (eBioscience), PE Rat Anti-Mouse CD25 (BD Biosciences), anti-mouse/rat FOXP3 APC (eBioscience), PE-Cy™7 Hamster Anti-Mouse CD11c (BD Biosciences), BB515 Rat Anti-Mouse I-A/I-E (BD Biosciences), PE Hamster Anti-Mouse CD80 (BD Biosciences), APC Rat Anti-Mouse CD49b (BD Biosciences), Ms CD69 PE H1.2F3 (BD Biosciences), FITC anti-mouse CD206 (MMR), PE Rat IgG1 κ Isotype Control RUO (BD Biosciences), Rat IgG2a kappa Isotype Control (eBR2a), PE (eBioscience), PE Rat IgG2a λ Isotype Control RUO (BD Biosciences), RAT IGG2A K ISO CNTL (EBR2A) APC (eBioscience), Ms CD274 BV650 MIH5 (BD Biosciences) and FITC anti-mouse H-2Kd/H-2Dd (Biolegend).

### RNA-seq

About 30 mg of fresh tumor tissues were placed in a centrifuge tube, mixed with 0.6 ml Trizol and ground with pre-cooled magnetic beads. 0.2 ml of chloroform was added to the homogenate. The mixed solution was shaken vigorously and left at room temperature for a few minutes and centrifuged. The upper aqueous phase was aspirated and mixed with anhydrous ethanol. Then the mixed solution was added to the adsorption column and centrifuged. The adsorption columns were washed twice with RPE solution and DEPC water was used to elute RNA. Transcriptome libraries were constructed with the kit VAHTS mRNA-seq V3 Library Prep Kit for Illumina (Vazyme). The transcriptome library sequencing was done by Oebiotech.

Based on the RNA-seq results, we analyzed differential genes, immune-related signature gene levels, T-cell infiltration levels, antigen presentation-related genes and cytotoxic molecule gene expression. The signature genes and analysis methods refer to published literature (*49, 50*). Differential expression was evaluated with DESeq. For a given gene to be considered significantly differentially expressed, we required a fold change of 1.5:1 or greater and a false discovery rate (FDR) corrected p-value of 0.05 or less. The following groups were compared: combination vs. mFc, Nb16 vs. mFc, SZU-101 vs. mFc. Differential gene expression was visualized by heatmap. Genes in the heatmap were clustered according to the log2(fold-change).

Based on the published literature (49, 50), immune signature scores are defined as the mean log2(fold-change) among all genes in each gene signature during immune-related signature gene levels analysis. We analyzed T cell infiltration within tumor tissues using ImmuCellAI (51).

### NDC safety analysis

Clinical observations were conducted daily from the day next to the randomization to experimental completion. At the end of experiment, mice were sacrificed. Blood and whole tissues and organs were collected. Animals were deeply anesthetized with isoflurane, followed by blood sample collection, and then euthanized by exsanguinated via the abdominal arteries or veins.

After isolation to obtain the serum, plasma chemistry was measured by biochemical analyzer (Cobas C501, Roche). Test indicators include alanine aminotransferase, aspartate aminotransferase, gamma glutamyl transferase, alkaline phosphatase, creatine kinase, total bilirubin, urea, creatinine, glucose, total cholesterol, triglycerides, total protein, albumin, globulin, albumin/globulin ratio, chloride, potassium and sodium. Alanine aminotransferase, aspartate aminotransferase, gamma glutamyl transferase, and alkaline phosphatase were detected by optimized IFCC method. Creatine kinase were detected by UV-test. Total bilirubin was detected by colorimetric diazomethod. Urea was detected by kinetic UV method with urease. Creatinine was detected by enzymatic colorimetry assay. Glucose was detected by enzymatic method with hexokinase. Total cholesterol and triglycerides were detected by enzymatic colorimetric assay. Total protein was detected by colorimetric assay with biuret. Albumin was detected by colorimetric assay with BCG. Chloride, potassium and sodium were detected by indirect ion selective electrode assay.

Hematology was measured by blood counters (Hemavet 950 FS). Test indicators include white blood cell (leukocyte) count, red blood cell (erythrocyte) count, hemoglobin, hematocrit, mean corpuscular (erythrocyte) volume, mean corpuscular (erythrocyte) hemoglobin, mean corpuscular (erythrocyte) hemoglobin concentration, red cell (erythrocyte volume) distribution width, platelet (thrombocyte) count, mean platelet (thrombocyte) volume, neutrophils count, percent of neutrophils, lymphocytes count, percent of lymphocytes, eosinophils count, percent of eosinophils, monocytes count, percent of monocytes, basophils count and percent of basophils.

Necropsy will be performed at the end of dosing phase. External features and orifices, the cranial cavity and external surfaces of the brain, and the thoracic, abdominal and pelvic cavities and associated organs/tissues and administration site will be completely examined during necropsy. The major organs of mice will be collected at necropsy for the animals from main study groups. After fixation, all preserved tissues/organs will be trimmed, dehydrated, embedded, sectioned, stained with hematoxylin and eosin (H&E) and then examined microscopically. Peer review of histopathology will be performed by a qualified pathologist.

Data file S1. Hematology data of 4G11-SZU-101.

Data file S2. Plasma chemistry data of 4G11-SZU-101.

Data file S3. Pathology analysis of 4G11-SZU-101.

---

**Fig. S1.**
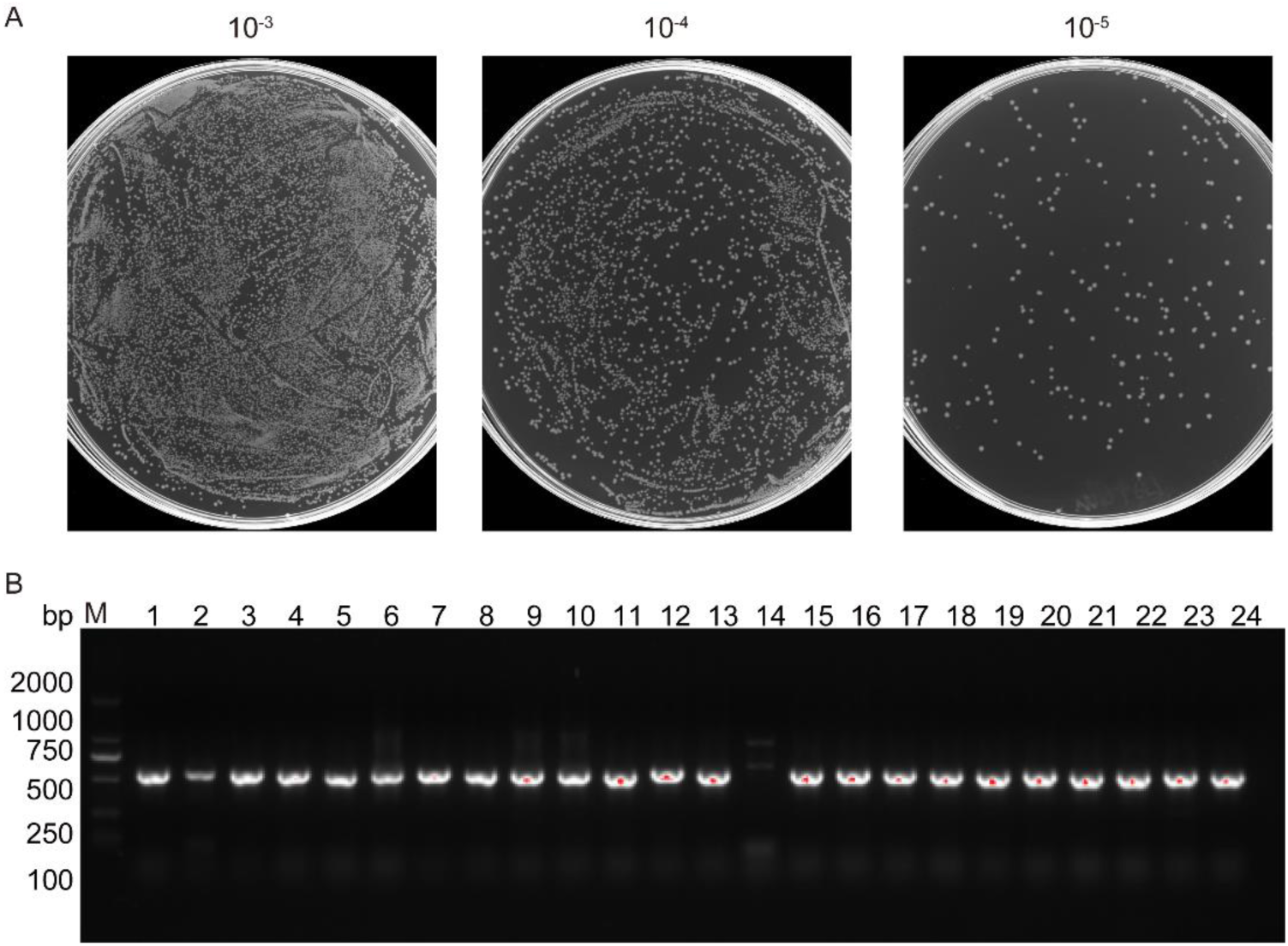
Nanobody library construction. (**A**) The library size was calculated by counting the colony numbers after gradient dilution. (**B**) The insertion rate was determined by picking 24 colonies randomly through colony PCR.

**Fig. S2.**
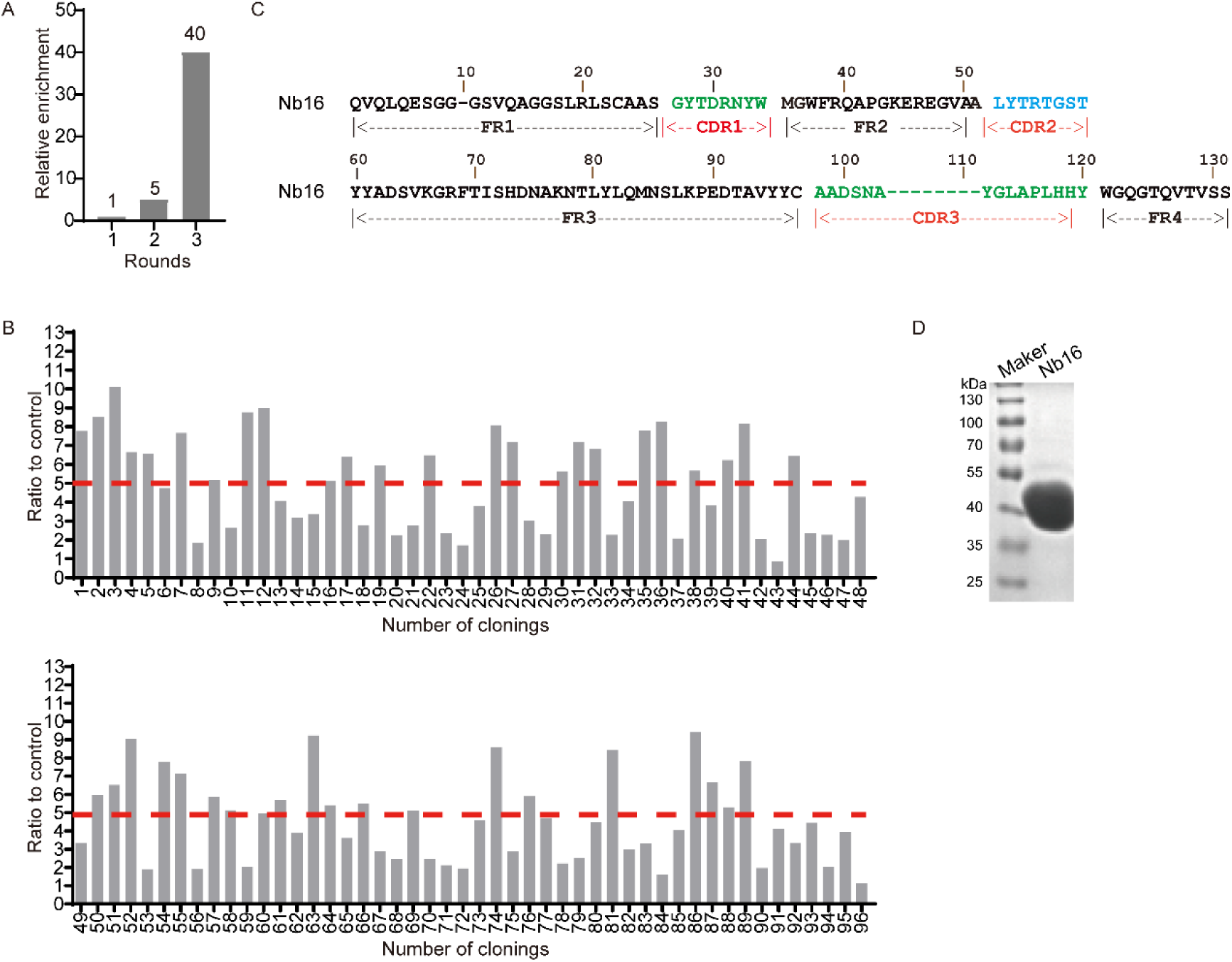
Phage library screening and anti-PD-L1 nanobody identification. (**A**) The relative enrichment ratio (40-fold) of anti-mPD-L1 nanobodies after 3 rounds of bio-panning. (**B**) 43 positive colonies were identified by PE-ELISA. Positive colonies had binding ratio higher than 5 relative to negative control. (**C**) The amino acid sequence of Nb16. (**D**) The SDS-PAGE was used to characterize the nanobodies (15 kDa) with an Fc tag (27 kDa).

**Fig. S3.**
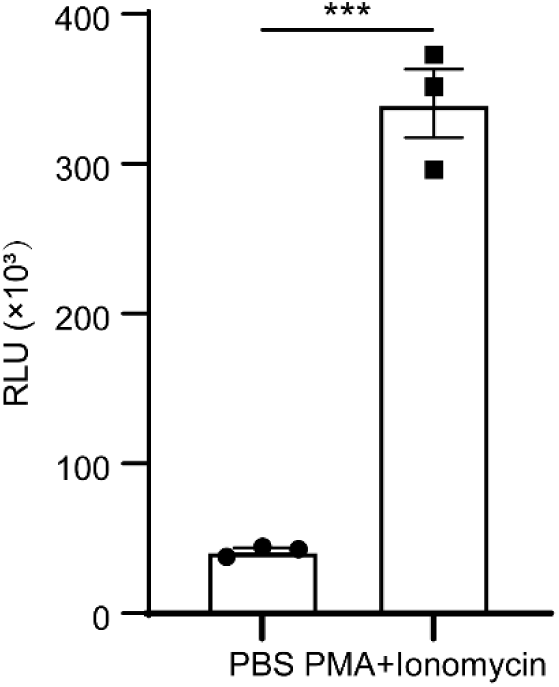
Construction of Jurkat-NFAT-luciferase cells. Jurkat-NFAT-luciferase cells were stimulated with 50 ng/mL of PMA and 1 µg/mL of ionomycin or PBS for 18 h. The RLU of luciferase expressed by Jurkat-NFAT-luciferase cells was measured by the automatic microplate reader SpectraMax. Data indicate mean ± SEM and are representative of at least two independent experiments. *** p < 0.001 as determined by unpaired t test.

**Fig. S4.**
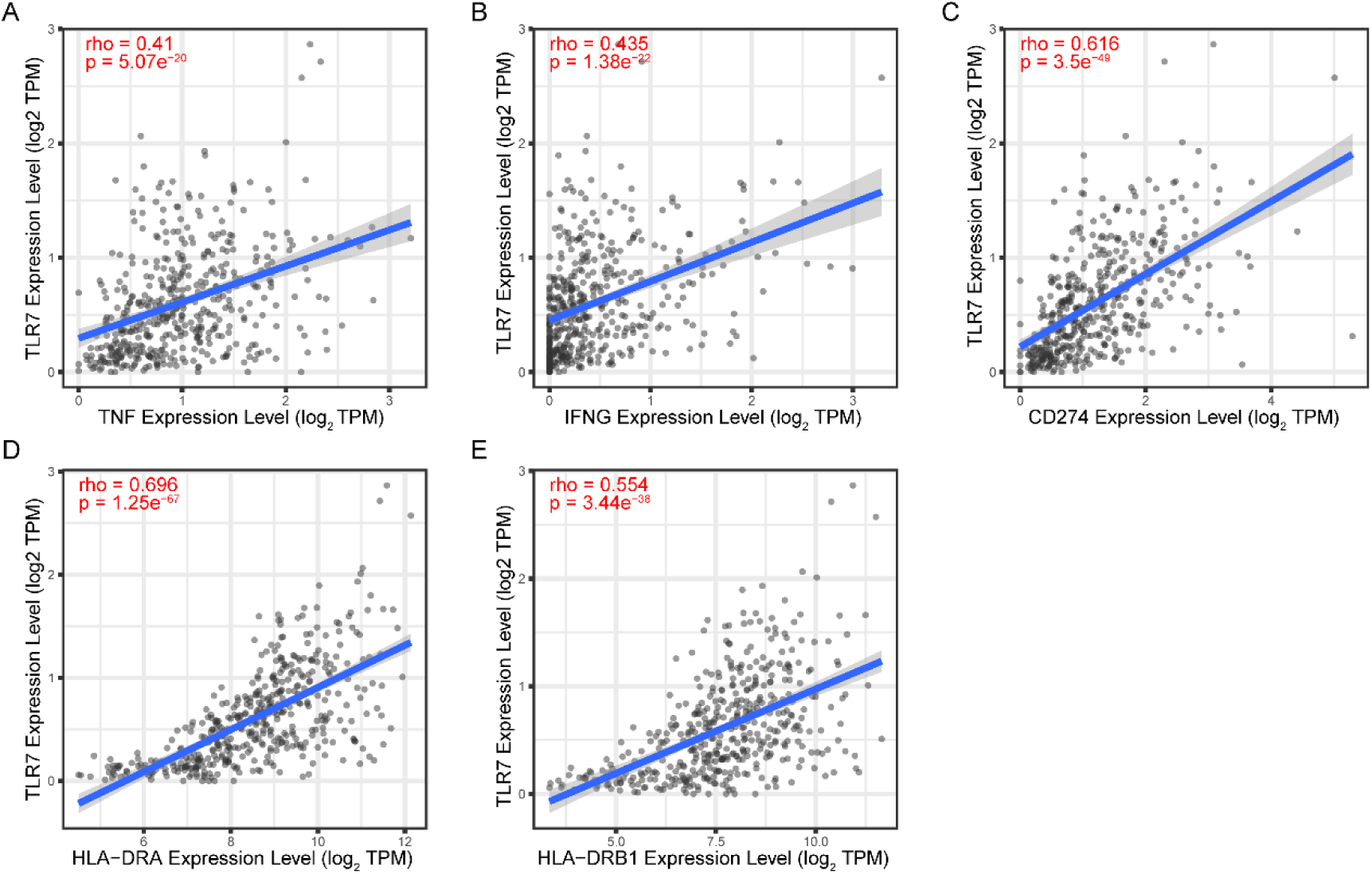
The expression association of TLR7 with TNF, IFNG, CD274, HLA-DRA and HLA-DRB1 in COAD cohorts. Correlation analysis between TLR7 and TNF (**A**), IFNG (**B**), CD274 (**C**), HLA-DRA (**D**) and HLA-DRB1 (**E**) in COAD from TCGA database were determined by TIMER2.0 and showed by scatter plots. Spearman’s rank correlation coefficient is represented by rho.

**Fig. S5.**
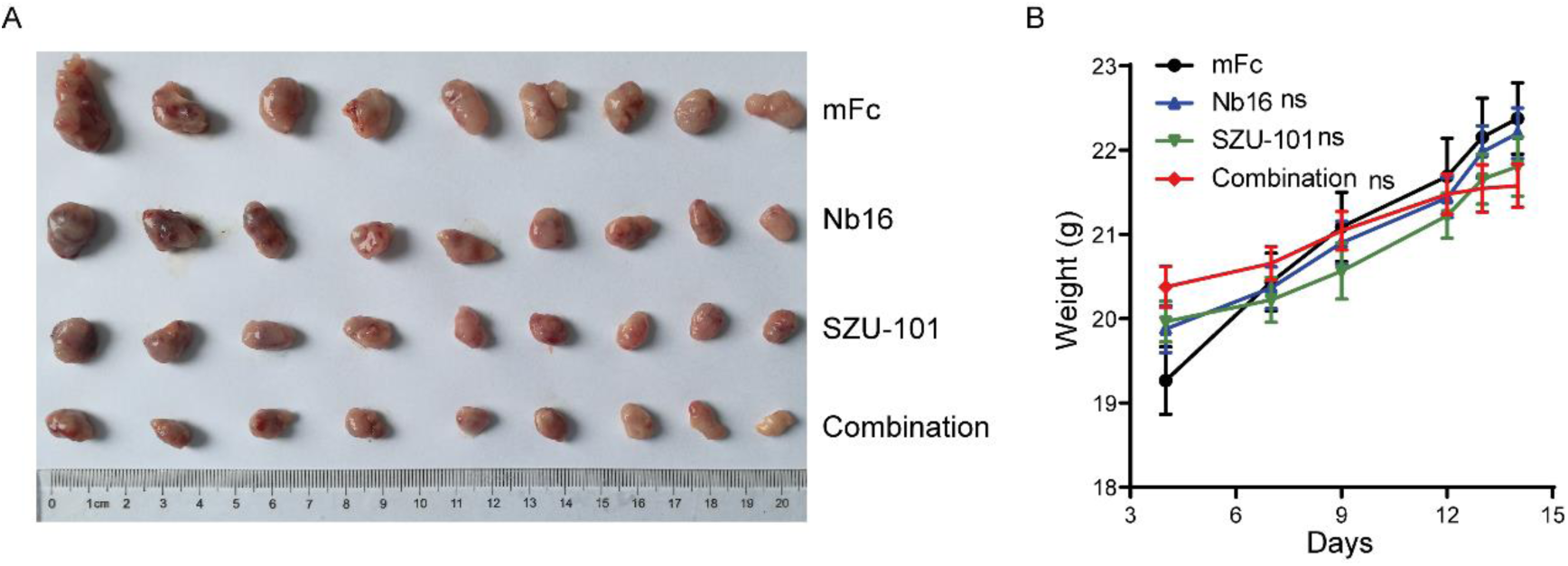
PD-L1 nanobody and TLR7 agonist exert synergistic antitumor effects. BALB/c mice were subcutaneously inoculated with CT26 cells (1×10^6^, induced by IFN-γ). Mice bearing tumors were treated with 200μg of mFc or 3 mg/kg of SZU-101 or 200 μg of Nb16 or combination. (**A**) The photograph shows the size of the tumor at the end of the experiment. (**B**) During the treatment, the weight changes of the mice were recorded and made into a curve plot. Data indicate mean ± SEM. ns not significant as determined by one-way ANOVA with multiple comparison tests.

**Fig. S6.**
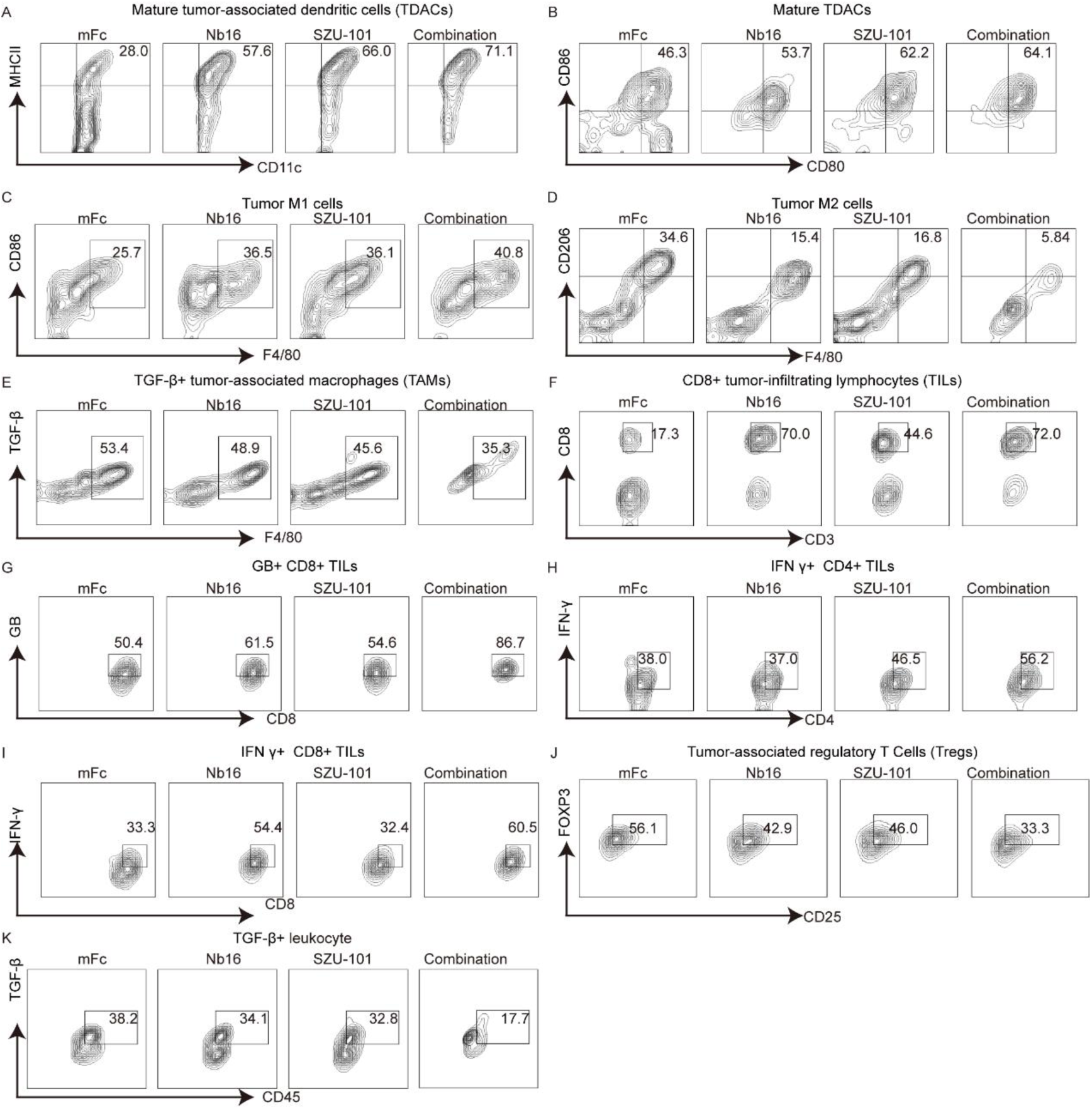
Immunophenotyping for Nb16 and SZU-101 combination therapy. **(A to K**) Immune effector cells in tumors treated with mFc, Nb16, SZU-101 or combination were quantified by flow cytometry (n = 3). Contour maps show the proportion of cells in a specific cell population. The percentages of MHC II^+^ DCs (A), mature DCs (B), M1 macrophages (C), M2 macrophages (D), TGF-β^+^ macrophages (E), CD8^+^ T cells (F), GB^+^ CD8+ T cells (G), IFN-γ^+^ CD4+ T cells (H), IFN-γ^+^ CD8+ T cells (I), Tregs (J) and TGF-β^+^ leukocytes (K) are shown based on their respective markers.

**Fig. S7.**
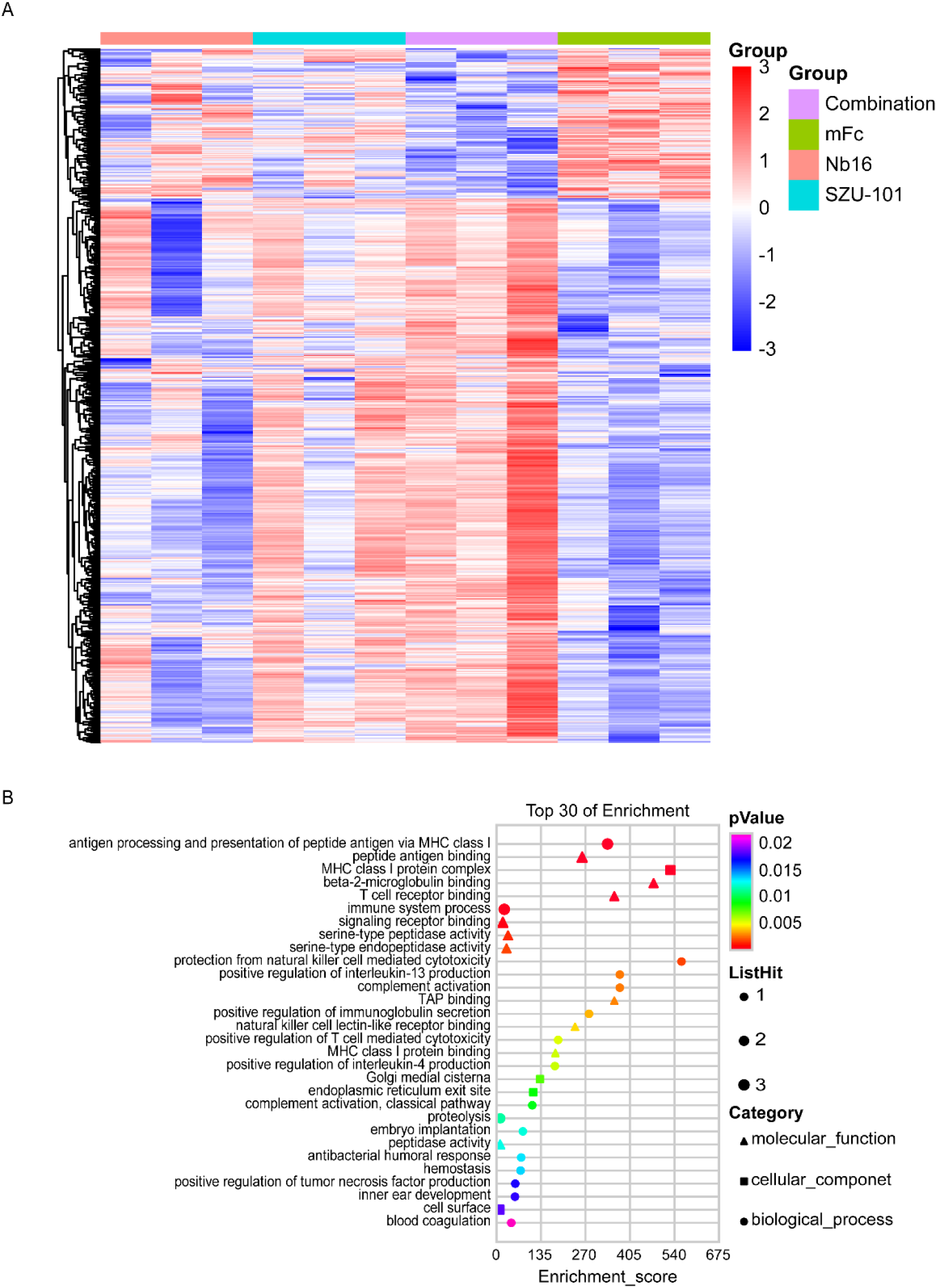
RNA-seq results for Nb16 and SZU-101 combination therapy. (**A**) Heatmap shows the gene expression for significantly differentially expressed genes from RNA-seq analysis. The colors in each box represent the log2(fold-change) in the expression of a gene in Nb16, SZU-101 or combination group relative to the median mFc group; rows represent individual genes, and columns represent individual mice. (**B**) The bubble plot shows the GO enrichment analysis of differential genes upregulated by combination treatment compared to mFc.

**Fig. S8.**
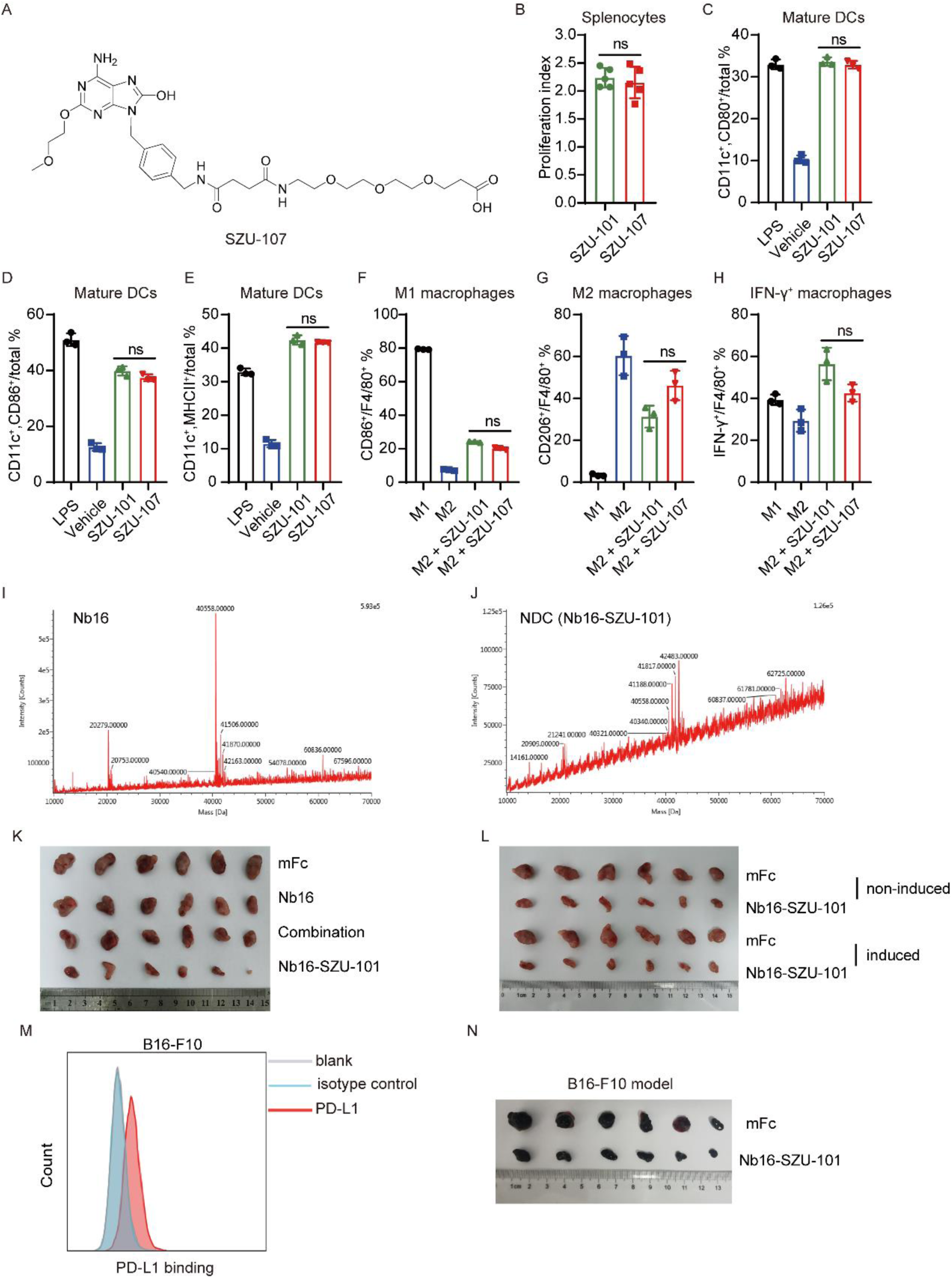
Identification of the PD-L1 and TLR7 dual-targeting nanobody-drug conjugate. (**A**) The structure of SZU-107. (**B**) Spleen cells obtained from BALB/c mice were treated with either SZU-101 or SZU-107 (50 μM, n = 5). An MTT assay was performed to determine the cell proliferation activity, shown as the proliferation index, which is calculated as the fold change with respect to the unstimulated control cultures. (**C** to **E**) BALB/c mouse bone marrow-derived dendritic cells (BMDCs) were treated with complete medium, LPS, 50 μM of SZU-101 or 50 μM of SZU-107 for 24 h (n = 3). CD80 (C), CD86 (D) and MHC II (E) on DCs were detected by flow cytometry. (**F** to **H**) BALB/c mouse bone marrow-derived macrophages (BMDMs) induced to differentiate into M2 macrophages were stimulated with 50 μM of SZU- 101 or 50 μM of SZU-107 for 24 h (n = 3). CD86 (F), CD206 (G) and IFN-γ (H) expression in macrophages was determined by flow cytometry. (**I** and **J**) Nb16 (I) and NDC (J) were identified by mass spectrometry, and the DAR of NDC was determined according to the change in the molecular weight of the antibody.(**K**) BALB/c mice were subcutaneously inoculated with 1×10^6^ of CT26 cells (induced by IFN-γ). Mice bearing tumors were treated with 200 μg of mFc, 200 μg of Nb16, combination or 200 μg of NDC. The photograph shows the size of the tumor at the end of the experiment. (**L**) BALB/c mice were subcutaneously inoculated with 1×10^6^ of CT26 cells (induced by IFN-γ or not). Mice bearing tumors were treated with 200 μg of mFc or 200 μg of NDC. The photograph shows the size of the tumor at the end of the experiment. (**M**) Flow cytometry histograms showing the expression levels of PD-L1 in B16-F10 cells. (**N**) C57BL/6 mice were subcutaneously inoculated with 1×10^6^ of B16-F10 cells. Mice bearing tumors were treated with 200 μg of mFc or 200 μg of NDC. The photograph shows the size of the tumor at the end of the experiment. Data indicate mean ± SEM. ns not significant as determined by one-way ANOVA with multiple comparison tests or unpaired t test.

**Fig. S9.**
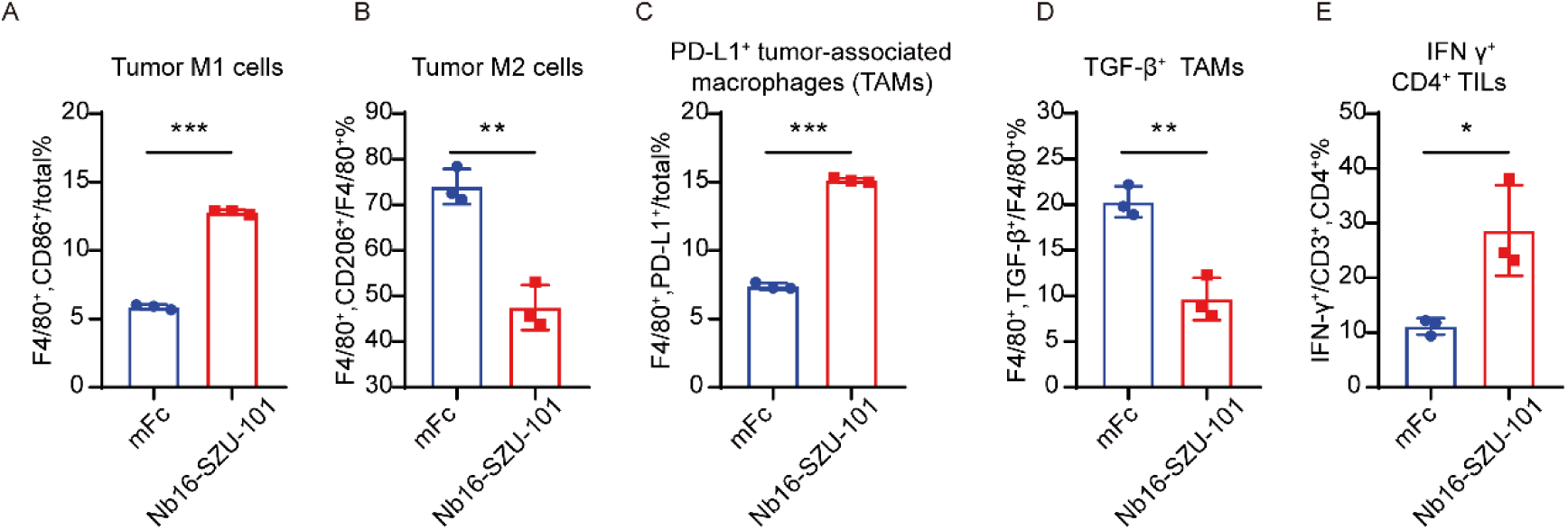
Immunophenotyping for NDC therapy. (**A** to **E**) Immune effector cells in tumors from mice treated with mFc or NDC were quantified by flow cytometry (n = 3). Scatter plots show the proportion of cells in a specific cell population. The percentages of M1 macrophages (A), M2 macrophages (B), PD-L1^+^ macrophages (C), TGF-β^+^ macrophages (D), IFN-γ^+^ CD4^+^ T cells (E) are shown based on their respective markers. The data are presented the mean ± SEM and are representative of at least two independent experiments. * p < 0.05; ** p < 0.01; *** p < 0.001; ns, not significant as determined by unpaired t test.

**Fig. S10.**
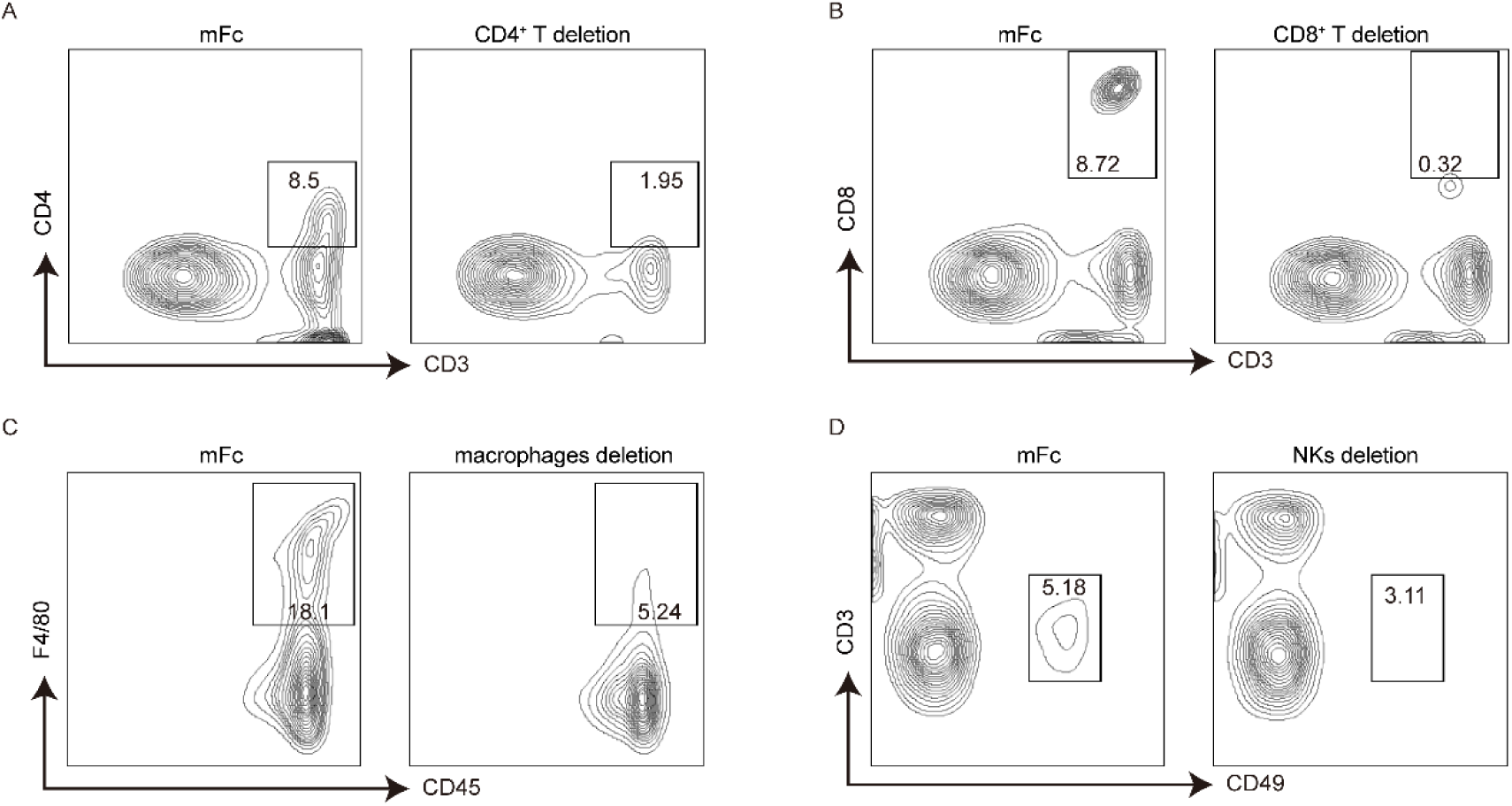
Validation of Immune cell deletion. (**A** to **D**) BALB/c mice were subcutaneously inoculated with 1×106 of CT26 cells (induced by IFN-γ) and treated with 200 μg of mFc or 200 μg of Nb16-SZU-101 dosed with anti-murine CD4, anti-murine CD8, anti-murine NK1.1, or clodronate liposomes. Deletion of specific cell populations was identified by flow cytometry. Contour maps show the proportion of cells in a specific cell population. The percentages of CD4^+^ T cells (A), CD8^+^ T cells (B), macrophages (C) and NK cells (D) are shown based on their respective markers.

**Fig. S11.**
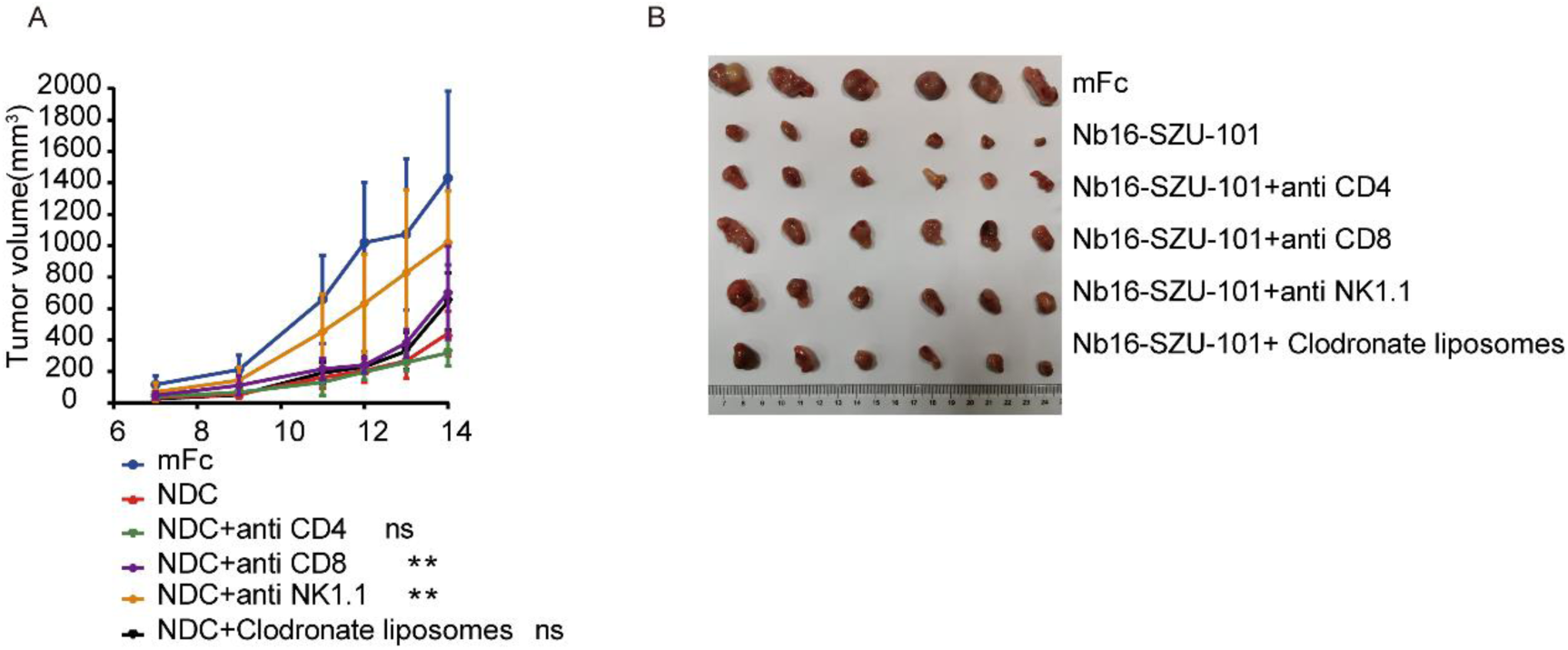
Immune cell deletion study. (**A** and **B**) BALB/c mice were subcutaneously inoculated with 1×10^6^ of CT26 cells (induced by IFN-γ) and treated with 200 μg of mFc or 200 μg of Nb16-SZU-101 dosed with anti-murine CD4, anti-murine CD8, anti-murine NK1.1, or clodronate liposomes. Tumor growth (A) in mice and tumor photos (B) are shown (n = 6). The data are presented as the mean ± SEM. ** p < 0.01; ns, not significant as determined by one-way ANOVA with multiple comparison tests.

**Fig. S12.**
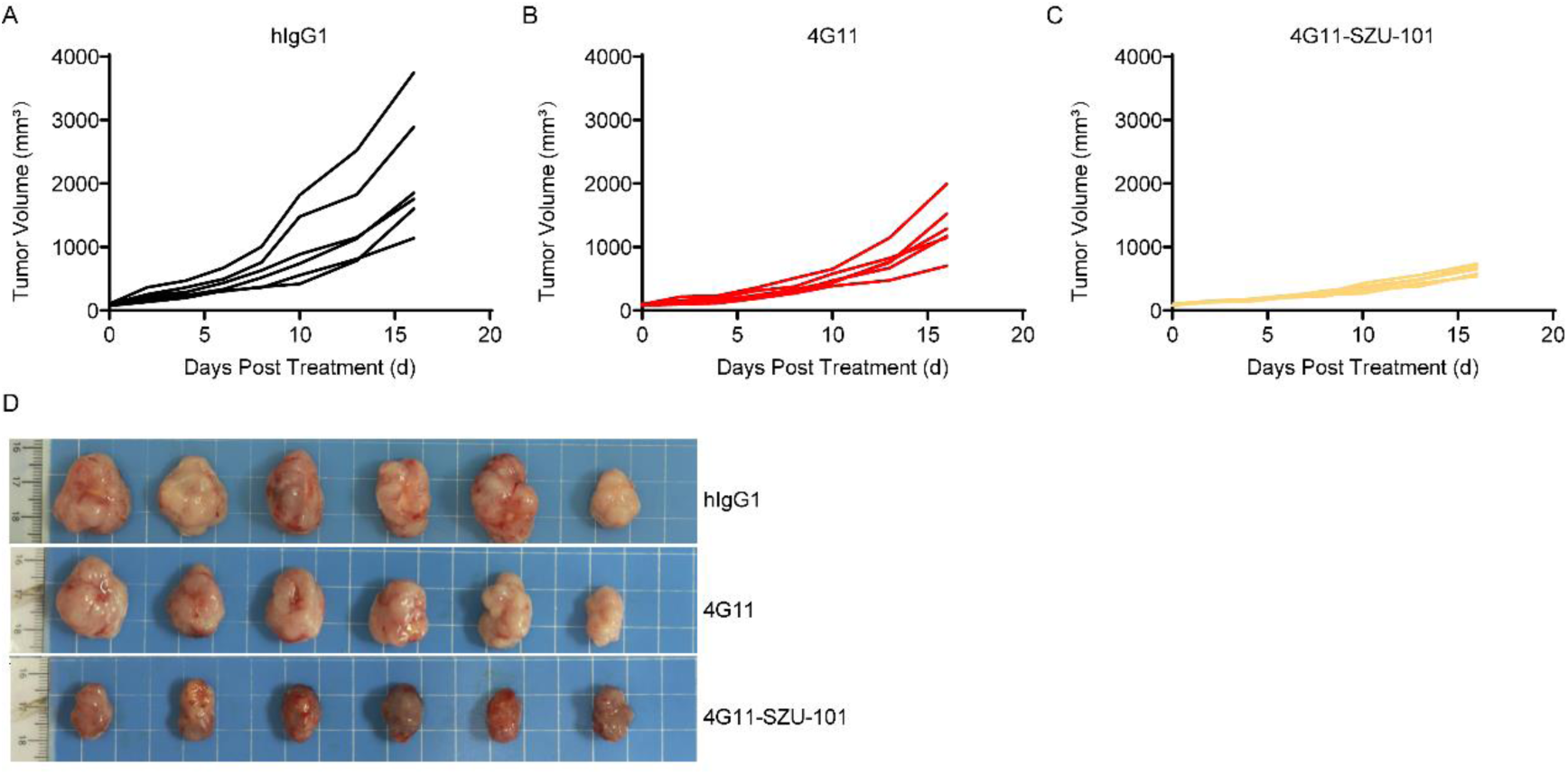
Antitumor activity of 4G11-SZU-101. PD-1/PD-L1 dual-humanized BALB/c mice were subcutaneously inoculated with CT26/hPD-L1 cells (5×10^5^). Mice bearing tumors were treated with 130 μg of hIgG1, 200 μg of 4G11 or 200 μg of 4G11-SZU-101. Tumor growth (**A** to **C**) in individual mice is shown ( n = 6). (**D**) The photograph shows the size of the tumor at the end of the experiment.

## References and Notes

1. H. Dong, G. Zhu, K. Tamada, L. Chen, B7-H1, a third member of the B7 family, co-stimulates T-cell proliferation and interleukin-10 secretion. Nature medicine. 5, 1365–1369 (1999).

2. H. Dong, S. E. Strome, D. R. Salomao, H. Tamura, F. Hirano, D. B. Flies, P. C. Roche, J. Lu, G. Zhu, K. Tamada, V. A. Lennon, E. Celis, L. Chen, Tumor-associated B7-H1 promotes T-cell apoptosis: a potential mechanism of immune evasion. Nat Med. 8, 793–800 (2002).

3. E. J. Wherry, T cell exhaustion. Nat Immunol. 12, 492–499 (2011).

4. J. R. Brahmer, S. S. Tykodi, L. Q. M. Chow, W.-J. Hwu, S. L. Topalian, P. Hwu, C. G. Drake, L. H. Camacho, J. Kauh, K. Odunsi, H. C. Pitot, O. Hamid, S. Bhatia, R. Martins, K. Eaton, S. Chen, T. M. Salay, S. Alaparthy, J. F. Grosso, A. J. Korman, S. M. Parker, S. Agrawal, S. M. Goldberg, D. M. Pardoll, A. Gupta, J. M. Wigginton, Safety and Activity of Anti–PD-L1 Antibody in Patients with Advanced Cancer. 366, 2455–2465 (2012).

5. H. Borghaei, L. Paz-Ares, L. Horn, D. R. Spigel, M. Steins, N. E. Ready, L. Q. Chow, E. E. Vokes, E. Felip, E. Holgado, F. Barlesi, M. Kohlhäufl, O. Arrieta, M. A. Burgio, J. Fayette, H. Lena, E. Poddubskaya, D. E. Gerber, S. N. Gettinger, C. M. Rudin, N. Rizvi, L. Crinò, G. R. Blumenschein, Jr., S. J. Antonia, C. Dorange, C. T. Harbison, F. Graf Finckenstein, J. R. Brahmer, Nivolumab versus Docetaxel in Advanced Nonsquamous Non-Small-Cell Lung Cancer. N Engl J Med. 373, 1627–1639 (2015).

6. R. L. Ferris, G. Blumenschein, Jr., J. Fayette, J. Guigay, A. D. Colevas, L. Licitra, K. Harrington, S. Kasper, E. E. Vokes, C. Even, F. Worden, N. F. Saba, L. C. Iglesias Docampo, R. Haddad, T. Rordorf, N. Kiyota, M. Tahara, M. Monga, M. Lynch, W. J. Geese, J. Kopit, J. W. Shaw, M. L. Gillison, Nivolumab for Recurrent Squamous-Cell Carcinoma of the Head and Neck. N Engl J Med. 375, 1856–1867 (2016).

7. R. J. Motzer, B. Escudier, D. F. McDermott, S. George, H. J. Hammers, S. Srinivas, S. S. Tykodi, J. A. Sosman, G. Procopio, E. R. Plimack, D. Castellano, T. K. Choueiri, H. Gurney, F. Donskov, P. Bono, J. Wagstaff, T. C. Gauler, T. Ueda, Y. Tomita, F. A. Schutz, C. Kollmannsberger, J. Larkin, A. Ravaud, J. S. Simon, L. A. Xu, I. M. Waxman, P. Sharma, Nivolumab versus Everolimus in Advanced Renal-Cell Carcinoma. N Engl J Med. 373, 1803–1813 (2015).

8. H. L. Kaufman, J. Russell, O. Hamid, S. Bhatia, P. Terheyden, S. P. D’Angelo, K. C. Shih, C. Lebbé, G. P. Linette, M. Milella, I. Brownell, K. D. Lewis, J. H. Lorch, K. Chin, L. Mahnke, A. von Heydebreck, J. M. Cuillerot, P. Nghiem, Avelumab in patients with chemotherapy-refractory metastatic Merkel cell carcinoma: a multicentre, single-group, open-label, phase 2 trial. The Lancet. Oncology. 17, 1374–1385 (2016).

9. R. S. Herbst, P. Baas, D. W. Kim, E. Felip, J. L. Pérez-Gracia, J. Y. Han, J. Molina, J. H. Kim, C. D. Arvis, M. J. Ahn, M. Majem, M. J. Fidler, G. de Castro, Jr., M. Garrido, G. M. Lubiniecki, Y. Shentu, E. Im, M. Dolled-Filhart, E. B. Garon, Pembrolizumab versus docetaxel for previously treated, PD-L1-positive, advanced non-small-cell lung cancer (KEYNOTE-010): a randomised controlled trial. Lancet (London, England). 387, 1540–1550 (2016).

10. X. Meng, Y. Liu, J. Zhang, F. Teng, L. Xing, J. Yu, PD-1/PD-L1 checkpoint blockades in non-small cell lung cancer: New development and challenges. Cancer Lett. 405, 29–37 (2017).

11. P. S. Hegde, D. S. Chen, Top 10 Challenges in Cancer Immunotherapy. Immunity. 52, 17–35 (2020).

12. J. Xin Yu, J. P. Hodge, C. Oliva, S. T. Neftelinov, V. M. Hubbard-Lucey, J. Tang, Trends in clinical development for PD-1/PD-L1 inhibitors. Nat Rev Drug Discov. 19, 163–164 (2020).

13. S. Upadhaya, S. T. Neftelino, J. P. Hodge, C. Oliva, J. R. Campbell, J. X. Yu, Combinations take centre stage in PD1/PDL1 inhibitor clinical trials. Nat Rev Drug Discov. (2020).

14. J. Haanen, Converting Cold into Hot Tumors by Combining Immunotherapies. Cell. 170, 1055–1056 (2017).

15. M. Binnewies, E. W. Roberts, K. Kersten, V. Chan, D. F. Fearon, M. Merad, L. M. Coussens, D. I. Gabrilovich, S. Ostrand-Rosenberg, C. C. Hedrick, R. H. Vonderheide, M. J. Pittet, R. K. Jain, W. Zou, T. K. Howcroft, E. C. Woodhouse, R. A. Weinberg, M. F. Krummel, Understanding the tumor immune microenvironment (TIME) for effective therapy. Nature Medicine. 24, 541–550 (2018).

16. J. C. Waite, B. Wang, L. Haber, A. Hermann, E. Ullman, X. Ye, D. Dudgeon, R. Slim, D. K. Ajithdoss, S. J. Godin, I. Ramos, Q. Wu, E. Oswald, P. Poon, J. Golubov, D. Grote, J. Stella, A. Pawashe, J. Finney, E. Herlihy, H. Ahmed, V. Kamat, A. Dorvilliers, E. Navarro, J. Xiao, J. Kim, S. N. Yang, J. Warsaw, C. Lett, L. Canova, T. Schulenburg, R. Foster, P. Krueger, E. Garnova, A. Rafique, R. Babb, G. Chen, N. Stokes Oristian, C. J. Siao, C. Daly, C. Gurer, J. Martin, L. Macdonald, D. MacDonald, W. Poueymirou, E. Smith, I. Lowy, G. Thurston, W. Olson, J. C. Lin, M. A. Sleeman, G. D. Yancopoulos, A. J. Murphy, D. Skokos, Tumor-targeted CD28 bispecific antibodies enhance the antitumor efficacy of PD-1 immunotherapy. Science translational medicine. 12, (2020).

17. Y. Liang, H. Tang, J. Guo, X. Qiu, Z. Yang, Z. Ren, Z. Sun, Y. Bian, L. Xu, H. Xu, J. Shen, Y. Han, H. Dong, H. Peng, Y. X. Fu, Targeting IFNalpha to tumor by anti-PD-L1 creates feedforward antitumor responses to overcome checkpoint blockade resistance. Nat Commun. 9, 4586 (2018).

18. X. Liu, X. Bao, M. Hu, H. Chang, M. Jiao, J. Cheng, L. Xie, Q. Huang, F. Li, C. Y. Li, Inhibition of PCSK9 potentiates immune checkpoint therapy for cancer. Nature. (2020).

19. S. S. Diebold, T. Kaisho, H. Hemmi, S. Akira, C. Reis e Sousa, Innate antiviral responses by means of TLR7-mediated recognition of single-stranded RNA. Science. 303, 1529–1531 (2004).

20. G. Trinchieri, A. Sher, Cooperation of Toll-like receptor signals in innate immune defence. Nature Reviews Immunology. 7, 179–190 (2007).

21. J. K. Geisse, P. Rich, A. Pandya, K. Gross, K. Andres, A. Ginkel, M. Owens, Imiquimod 5% cream for the treatment of superficial basal cell carcinoma: a double-blind, randomized, vehicle-controlled study. J Am Acad Dermatol. 47, 390–398 (2002).

22. K. A. Michaelis, M. A. Norgard, X. Zhu, P. R. Levasseur, S. Sivagnanam, S. M. Liudahl, K. G. Burfeind, B. Olson, K. R. Pelz, D. M. Angeles Ramos, H. C. Maurer, K. P. Olive, L. M. Coussens, T. K. Morgan, D. L. Marks, The TLR7/8 agonist R848 remodels tumor and host responses to promote survival in pancreatic cancer. Nat Commun. 10, 4682 (2019).

23. J. Zhu, S. He, J. Du, Z. Wang, W. Li, X. Chen, W. Jiang, D. Zheng, G. Jin, Local administration of a novel Toll-like receptor 7 agonist in combination with doxorubicin induces durable tumouricidal effects in a murine model of T cell lymphoma. J Hematol Oncol. 8, 21 (2015).

24. M. Singh, H. Khong, Z. Dai, X. F. Huang, J. A. Wargo, Z. A. Cooper, J. P. Vasilakos, P. Hwu, W. W. Overwijk, Effective innate and adaptive antimelanoma immunity through localized TLR7/8 activation. J Immunol. 193, 4722–4731 (2014).

25. M. P. Schon, M. Schon, TLR7 and TLR8 as targets in cancer therapy. Oncogene. 27, 190–199 (2008).

26. S. E. Ackerman, C. I. Pearson, J. D. Gregorio, J. C. Gonzalez, J. A. Kenkel, F. J. Hartmann, A. Luo, P. Y. Ho, H. LeBlanc, L. K. Blum, S. C. Kimmey, A. Luo, M. L. Nguyen, J. C. Paik, L. Y. Sheu, B. Ackerman, A. Lee, H. Li, J. Melrose, R. P. Laura, V. C. Ramani, K. A. Henning, D. Y. Jackson, B. S. Safina, G. Yonehiro, B. H. Devens, Y. Carmi, S. J. Chapin, S. C. Bendall, M. Kowanetz, D. Dornan, E. G. Engleman, M. N. Alonso, Immune-stimulating antibody conjugates elicit robust myeloid activation and durable antitumor immunity. Nature Cancer. (2020).

27. X. Wang, B. Yu, B. Cao, J. Zhou, Y. Deng, Z. Wang, G. Jin, A chemical conjugation of JQ-1 and a TLR7 agonist induces tumoricidal effects in a murine model of melanoma via enhanced immunomodulation. Int J Cancer. 148, 437–447 (2021).

28. V. Lombardi, L. Van Overtvelt, S. Horiot, P. Moingeon, Human dendritic cells stimulated via TLR7 and/or TLR8 induce the sequential production of Il-10, IFN-gamma, and IL-17A by naive CD4+ T cells. J Immunol. 182, 3372–3379 (2009).

29. S. Akira, K. Takeda, T. Kaisho, Toll-like receptors: critical proteins linking innate and acquired immunity. Nat Immunol. 2, 675–680 (2001).

30. D. Q. Wang, M. Precopio, T. Lan, D. Yu, J. X. Tang, E. R. Kandimalla, S. Agrawal, Antitumor Activity and Immune Response Induction of a Dual Agonist of Toll-Like Receptors 7 and 8. Mol Cancer Ther. 9, 1788–1797 (2010).

31. E. Koga-Yamakawa, S. J. Dovedi, M. Murata, H. Matsui, A. J. Leishman, J. Bell, D. Ferguson, S. P. Heaton, T. Oki, H. Tomizawa, A. Bahl, H. Takaku, R. W. Wilkinson, H. Harada, Intratracheal and oral administration of SM-276001: A selective TLR7 agonist, leads to antitumor efficacy in primary and metastatic models of cancer. International Journal of Cancer. 132, 580–590 (2013).

32. J. Z. Oh, J. S. Kurche, M. A. Burchill, R. M. Kedl, TLR7 enables cross-presentation by multiple dendritic cell subsets through a type I IFN-dependent pathway. Blood. 118, 3028–3038 (2011).

33. S. R. Mullins, J. P. Vasilakos, K. Deschler, I. Grigsby, P. Gillis, J. John, M. J. Elder, J. Swales, E. Timosenko, Z. Cooper, S. J. Dovedi, A. J. Leishman, N. Luheshi, J. Elvecrog, A. Tilahun, R. Goodwin, R. Herbst, M. A. Tomai, R. W. Wilkinson, Intratumoral immunotherapy with TLR7/8 agonist MEDI9197 modulates the tumor microenvironment leading to enhanced activity when combined with other immunotherapies. J Immunother Cancer. 7, 244 (2019).

34. E. Henke, R. Nandigama, S. Ergun, Extracellular Matrix in the Tumor Microenvironment and Its Impact on Cancer Therapy. Front Mol Biosci. 6, 160 (2019).

35. I. Van Audenhove, J. Gettemans, Nanobodies as Versatile Tools to Understand, Diagnose, Visualize and Treat Cancer. EBioMedicine. 8, 40–48 (2016).

36. M. Kijanka, B. Dorresteijn, S. Oliveira, P. M. P. V. E. Henegouwen, Nanobody-based cancer therapy of solid tumors. Nanomedicine-Uk. 10, 161–174 (2015).

37. E. Y. Yang, K. Shah, Nanobodies: Next Generation of Cancer Diagnostics and Therapeutics. Front Oncol. 10, 1182 (2020).

38. W. Yin, X. Yu, X. Kang, Y. Zhao, P. Zhao, H. Jin, X. Fu, Y. Wan, C. Peng, Y. Huang, Remodeling Tumor-Associated Macrophages and Neovascularization Overcomes EGFR(T790M) -Associated Drug Resistance by PD-L1 Nanobody-Mediated Codelivery. Small. 14, e1802372 (2018).

39. S. Muyldermans, Nanobodies: natural single-domain antibodies. Annu Rev Biochem. 82, 775–797 (2013).

40. F. Macian, NFAT proteins: Key regulators of T-cell development and function. Nature Reviews Immunology. 5, 472–484 (2005).

41. N. Eiro, L. Gonzalez, L. O. Gonzalez, A. Andicoechea, M. Fernandez-Diaz, A. Altadill, F. J. Vizoso, Study of the expression of toll-like receptors in different histological types of colorectal polyps and their relationship with colorectal cancer. J Clin Immunol. 32, 848–854 (2012).

42. J. H. Han, S. Y. Park, J. B. Kim, S. D. Cho, B. Kim, B. Y. Kim, M. J. Kang, D. J. Kim, J. H. Park, J. H. Park, TLR7 expression is decreased during tumour progression in transgenic adenocarcinoma of mouse prostate mice and its activation inhibits growth of prostate cancer cells. Am J Reprod Immunol. 70, 317–326 (2013).

43. J. Jiang, L. Dong, B. Qin, H. Shi, X. Guo, Y. Wang, Decreased expression of TLR7 in gastric cancer tissues and the effects of TLR7 activation on gastric cancer cells. Oncol Lett. 12, 631–636 (2016).

44. T. Li, J. Fu, Z. Zeng, D. Cohen, J. Li, Q. Chen, B. Li, X. S. Liu, TIMER2.0 for analysis of tumor-infiltrating immune cells. Nucleic Acids Res. 48, W509–W514 (2020).

45. M. Fabbri, A. Paone, F. Calore, R. Galli, E. Gaudio, R. Santhanam, F. Lovat, P. Fadda, C. Mao, G. J. Nuovo, N. Zanesi, M. Crawford, G. H. Ozer, D. Wernicke, H. Alder, M. A. Caligiuri, P. Nana-Sinkam, D. Perrotti, C. M. Croce, MicroRNAs bind to Toll-like receptors to induce prometastatic inflammatory response. Proc Natl Acad Sci U S A. 109, E2110–2116 (2012).

46. J. P. Pradere, D. H. Dapito, R. F. Schwabe, The Yin and Yang of Toll-like receptors in cancer. Oncogene. 33, 3485–3495 (2014).

47. M. Dajon, K. Iribarren, I. Cremer, Dual roles of TLR7 in the lung cancer microenvironment. OncoImmunology. 4, (2015).

48. A. Mantovani, S. Sozzani, M. Locati, P. Allavena, A. Sica, Macrophage polarization: tumor-associated macrophages as a paradigm for polarized M2 mononuclear phagocytes. Trends Immunol. 23, 549–555 (2002).

49. Y. Lan, D. Zhang, C. Xu, K. W. Hance, B. Marelli, J. Qi, H. Yu, G. Qin, A. Sircar, V. M. Hernández, M. H. Jenkins, R. E. Fontana, A. Deshpande, G. Locke, H. Sabzevari, L. Radvanyi, K. M. Lo, Enhanced preclinical antitumor activity of M7824, a bifunctional fusion protein simultaneously targeting PD-L1 and TGF-β. Science translational medicine. 10, (2018).

50. S. Nakao, Y. Arai, M. Tasaki, M. Yamashita, R. Murakami, T. Kawase, N. Amino, M. Nakatake, H. Kurosaki, M. Mori, M. Takeuchi, T. Nakamura, Intratumoral expression of IL-7 and IL-12 using an oncolytic virus increases systemic sensitivity to immune checkpoint blockade. Science translational medicine. 12, (2020).

51. Y. R. Miao, Q. Zhang, Q. Lei, M. Luo, G. Y. Xie, H. Wang, A. Y. Guo, ImmuCellAI: A Unique Method for Comprehensive T-Cell Subsets Abundance Prediction and its Application in Cancer Immunotherapy. Advanced science (Weinheim, Baden-Wurttemberg, Germany). 7, 1902880 (2020).

52. J. Galon, D. Bruni, Approaches to treat immune hot, altered and cold tumours with combination immunotherapies. Nat Rev Drug Discov. 18, 197–218 (2019).

53. M. Yi, D. Jiao, H. Xu, Q. Liu, W. Zhao, X. Han, K. Wu, Biomarkers for predicting efficacy of PD-1/PD-L1 inhibitors. Mol Cancer. 17, 129 (2018).

54. Z. R. Lu, P. Qiao, Drug Delivery in Cancer Therapy, Quo Vadis? Mol Pharm. 15, 3603–3616 (2018).

55. T. Chanier, P. Chames, Nanobody Engineering: Toward Next Generation Immunotherapies and Immunoimaging of Cancer. Antibodies (Basel). 8, (2019).

56. R. S. Herbst, J. C. Soria, M. Kowanetz, G. D. Fine, O. Hamid, M. S. Gordon, J. A. Sosman, D. F. McDermott, J. D. Powderly, S. N. Gettinger, H. E. Kohrt, L. Horn, D. P. Lawrence, S. Rost, M. Leabman, Y. Xiao, A. Mokatrin, H. Koeppen, P. S. Hegde, I. Mellman, D. S. Chen, F. S. Hodi, Predictive correlates of response to the anti-PD-L1 antibody MPDL3280A in cancer patients. Nature. 515, 563–567 (2014).

